# Frost and drought: effects of extreme weather events on stem carbon dynamics in a Mediterranean beech forest

**DOI:** 10.1101/806026

**Authors:** Ettore D’Andrea, Negar Rezaie, Peter Prislan, Jozica Gričar, Jan Muhr, Alessio Collalti, Giorgio Matteucci

## Abstract

The effects of short-term extreme events on tree functioning and physiology are still rather elusive. European beech is one of the most sensitive species to late frost and water shortage. We investigated the intra-annual C dynamics in stems under such conditions.

Wood formation and stem CO_2_ efflux were monitored in a Mediterranean beech forest for three years (2015–2017), including a late frost (2016) and a summer drought (2017).

The late frost reduced radial growth and, consequently, the amount of carbon fixed in the stem biomass by 80%. Stem carbon efflux in 2016 was reduced by 25%, which can be attributed to the reduction of effluxes due to growth respiration. Counter to our expectations, we found no effects of the 2017 summer drought on radial growth and stem carbon efflux.

The studied extreme weather events had various effects on tree growth. Even though late spring frost had a devastating impact on beech radial growth in the current year, trees fully recovered in the following growing season, indicating high resilience of beech to this stressful event.

## Introduction

Even small changes in the mean or variation of a climate variable cause disproportionally large changes in the frequency of extreme weather events, recognized as major drivers of current and future ecosystem dynamics (Frank *et al*. 2015). In the near future, the Mediterranean region is predicted to be the most vulnerable of the European regions to global change (Schröter *et al*. 2005). Changes in temperature and precipitation regimes may increase drought risk (Schröter *et al*. 2005), which can negatively affect physiological performance (Rezaie *et al*. 2018), as well as the growth and competitive strength (Peuke, Schraml, Hartung & Rennenberg 2002) of common beech, one of the most important and widespread broadleaved trees in Europe.

Increasing spring temperatures can trigger earlier leaf unfolding (Gordo & Sanz 2010; Allevato *et al*. 2019), which in turn results in a higher risk that young leaves are exposed to late spring frost (Augspurger 2013), especially at higher elevations (Vitasse, Schneider, Rixen, Christen & Rebetez 2018). Temperatures below −4°C can kill the developing new shoots and leaves, thus reducing the photosynthetic area and ultimately the trees’ growth. In the case of late frost, depending on the intensity of damage, the formation of new leaves requires a high amount of reserves (Dittmar, Fricke & Elling 2006; D’Andrea *et al*. 2019).

Tree stems play an important role in the carbon balance of forest ecosystems (Yang, He, Aubrey, Zhuang & Teskey 2016). Part of the carbon (C) fixed by photosynthesis is allocated to the stem, and some is respired by stems and emitted into the atmosphere. Radial growth – an often used proxy for the overall allocation of C to the stem (Bascietto, Cherubini & Scarascia-Mugnozza 2004; Cuny *et al*. 2015; Chan, Berninger, Kolari, Nikinmaa & Hölttä 2018) – is largely related to the process of wood formation, which can be divided into five (main) developmental phases: *i*) cambial cell division; *ii*) cell enlargement; *iii*) secondary wall deposition and, *iv*) cell wall thickening (lignification); while *v*) in the case of vessels and fibres, also genetically-programmed cell death or *apoptosis* (Prislan, Čufar, De Luis & Gričar 2018). The whole process is sensitive to many factors, such as leaf phenology (Michelot, Simard, Rathgeber, Dufrêne & Damesin 2012), temperature (Begum, Nakaba, Oribe, Kubo & Funada 2007), drought (Linares, Camarero & Carreira 2009), tree-size and social status (Rathgeber, Rossi & Bontemps 2011) and tree vigour (Gričar, Krže & Čufar 2009).

As also reported by Damesin (2002), stem respiration may represent up to 1/3 of the overall above ground respiration and 1.4% of the annual carbon assimilation. However, field measurements of actual stem respiration (RS) are difficult if not impossible (Teskey, Saveyn, Steppe & McGuire 2008), and the most commonly measured proxy, stem CO_2_ efflux (ES), is likely to underestimate local respiration (Trumbore, Angert, Kunert, Muhr & Chambers 2013). Previous studies have reported a strong correlation between RS and ES, with ES ranging between 82–94% and 86–91% of RS in *Populus deltoides* W.Bartram ex Marshall (Saveyn, Steppe, Mc Guire, Lemeur & Teskey 2008) and *Dacrydium cupressinum* Lamb stems (Bowman *et al*. 2005), respectively. Another study using O_2_ uptake, as an alternative proxy for actual respiration, and comparing it to traditionally measured ES, showed that ES on average underestimated RS by about 41% (CO_2_ was not emitted locally at the point of measurement) (Hilman *et al*. 2019). ES and RS are different because part of the CO_2_ produced by respiration is not released directly through the bark into the atmosphere, but is dissolved in xylem sap and is carried upward by the transpiration stream (Bloemen *et al*. 2014). In addition, ES is affected by CO_2_ deriving from root respiration, which is also carried upward into the stem (Bloemen, McGuire, Aubrey, Teskey & Steppe 2013). Moreover, part of respired CO_2_ can be fixed in xylem storage pools (Ubierna *et al*. 2009). A recent global estimate showed that the stem CO_2_ efflux (ES) alone, from boreal to tropical forests combined, was 6.7 (± 1.1) Pg C yr^−1^, accounting for 11% and 20% of global forest ecosystem gross primary production (GPP) and net primary production (NPP), respectively (Yang *et al*. 2016).

ES is influenced by many factors, such as air temperature (Yang et al., 2016), growth rate (Damesin, Ceschia, Le Goff, Ottorini & Dufrêne 2002), distribution and turnover of living cells (Collalti *et al*. 2019), nitrogen concentration (Ceschia, Damesin, Lebaube, Pontailler & Dufrêne 2002) and tree social class (Guidolotti, Rey, D’Andrea, Matteucci & De Angelis 2013). Both RS and ES are also affected by growth respiration (*R_G_*), which provides the energy for synthesizing new tissues; and by maintenance respiration (*R_M_*), which maintains existing living cells (Ceschia *et al*. 2002). Separating ES into these components allows further investigation of stem carbon budgeting and tissue costs (Chan *et al*. 2018).

Despite the crucial role of extreme events and increasing attention on their prospective increasing role in future climate scenarios, information on the effect of short-term extreme events on tree functioning is still fairly elusive (Carrer, Brunetti & Castagneri 2016; Gazol *et al*. 2019). In this respect, not much is known about the interaction between wood formation (xylogenesis) and ES. At the seasonal timescale, the capacity of micro-coring technique to identify phenological phases of wood formation allows to attribute metabolic costs to each one of them (Meir, Mencuccini & Coughlin 2019).

A deeper investigation of this link is crucial, especially in the context of climate change, associated with increased frequency of extreme weather events (e.g., drought and late frost), which may greatly modify the contribution of these processes to the C cycle.

In this context, we monitored xylogenesis, together with ES and overall growth, in a mature Mediterranean beech forest (*Fagus sylvatica* L.) from 2015 to 2017 – a period characterized by a spring late frost (2016) and a summer drought (2017) – with the objective of unravelling the intra-annual C dynamics in stems under different climatic conditions and in response to extreme weather events. On the hypothesis that extreme weather events would alter the stem C dynamics at both tree and stand scales, we investigated tree scale physiological processes; specifically, of growth (xylogenesis) and respiration (proxied by stem CO_2_ efflux), and then upscaled to stand-scale growth and C emission related to this process. We hypothesized that: 1) cambial activity and radial growth may have ceased soon after leaf death due to the 2016 spring late frost; 2) second leaf re-sprouting starts at the expense of stem growth; 3) the 2017 summer drought would have negatively impacted stem biomass production and effluxes; and that, 4) climatic variability and extreme weather events are important factors in C dynamics on tree and stand scales.

## Material and methods

### Study site and description of weather events

The measurements were carried out between 2015 and 2017 on a long-term monitored beech stand (*Fagus sylvatica* L.) located at Selva Piana (41°50’58” N, 13°35’17” E, 1,560 m elevation), close to Collelongo (Abruzzi Region, Italy) in the Central Apennines. The site, established in 1991 and since 2006 part of the long-term ecological research (LTER) network, is located in a 3000 ha forest included in the wider forest area of the external belt of Abruzzi National Park. In 2017, the stand density was 725 trees ha^−1^, the basal area was 45.77 m^2^ ha^−1^ with a mean diameter at breast height (DBH) of 28.5 cm, and a mean tree height of 23 m. Mean tree age in 2013 was estimated to be about 110 years. Site topography is gently sloping and the soil is humic alisol with a variable depth (40–100 cm), developed on calcareous bedrock (Chiti *et al*. 2010). The climate is of Mediterranean mountain type, during the period 1989 – 2014 the mean annual temperature was 7.2°C, and the mean annual precipitation was 1178 mm, of which ∼10% falls in summer (Guidolotti *et al*. 2013).

In the night between the 25^th^ and 26^th^ of April 2016 (Day of Year, DOY 115), a spring late frost occurred in Central and Southern Italy, causing leaf damage in many beech stands (Bascietto, Bajocco, Mazzenga & Matteucci 2018; Greco *et al*. 2018; Nolè, Rita, Ferrara & Borghetti 2018; Allevato *et al*. 2019). At the Selva Piana site, the air temperature reached – 6°C at canopy level, destroying the whole-stand canopy and leaving the trees without leaves for almost two months (Bascietto *et al*. 2018; D’Andrea *et al*. 2019).

In the summer of 2017, a widespread positive temperature anomaly affected Central Italy and the Balkans, with a duration ranging from 20 to 35 days (Rita *et al*. 2019). In the same year, annual precipitation was 950 mm, with only 54 mm of precipitation throughout the entire summer; while only 1 mm of rain and a mean air temperature (23.9 °C) ∼ 2°C warmer than the long-term average (1989–2014) characterized August 2017, leading to a Standardized Precipitation Evapotranspiration Index (SPEI) > 1.5 during the vegetative season (Fig. 1).

**Fig. 1:**
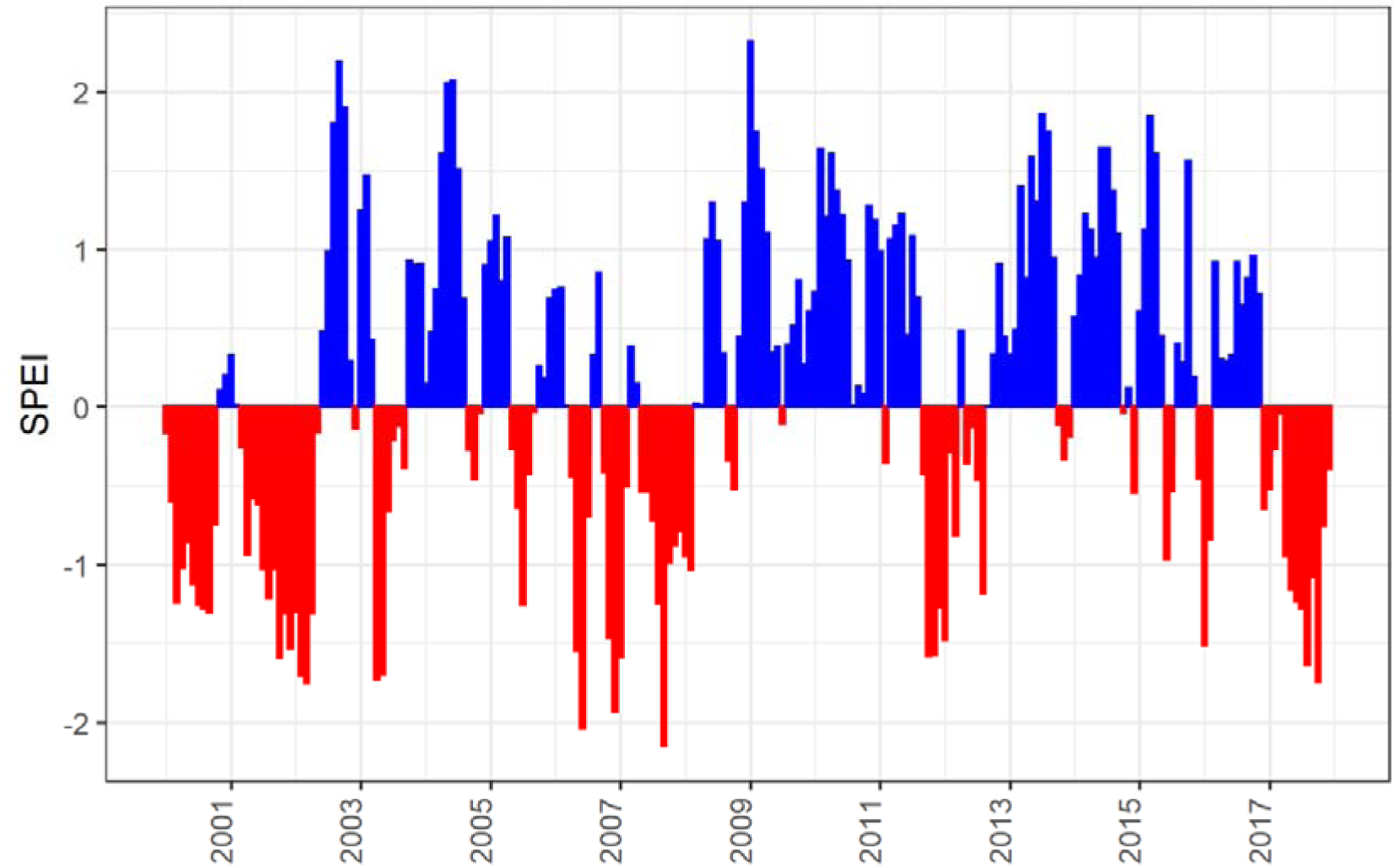
3-month Standardized Precipitation Evapotranspiration Index (SPEI) at the experimental site of Selva Piana-Collelongo.

### Tree selection, wood formation dynamics, xylem phenology and C fixation

Sampling was performed on five trees that were selected for their similarity, with a site tree ring chronology following the methodology described in Rezaie *et al*. (2018) and as done in other studies on wood formation and stem CO_2_ efflux (e.g. Ceschia *et al*., 2002; Damesin *et al*., 2002; Gruber *et al*., 2009; Delpierre *et al*., 2019). Microcore collection and ES measurements were carried out from April 2015, before leaf unfolding, until November 2017, when the trees had completely lost their leaves.

Microcores (2 mm diameter and 15 mm long) were extracted from each tree at 1.1-1.7 m above ground using a Trephor tool (Rossi, Menardi, Fontanella & Anfodillo 2005). Microcores were collected 15, 12, and 14 times in 2015, 2016, and 2017, respectively. To avoid wound effects, cores were sampled at a distance of at least 5 cm from each other. The microcores, containing bark, cambium, newly developing xylem and 1-2 older xylem rings, were immediately stored in formaldehyde-ethanol-acetic acid solution (FEA) in the field. Cross-sections of the microcores were prepared following the standard methodology (Prislan, Gričar, De Luis, Smith & Cufar 2013) and were photographed in high definition under a Leica DM 4000 microscope (Leica Microsystems, Wetzlar, Germany) using transmission and polarized light. Histometrical analyses were performed on images taken with a Leica DFC 280 digital camera using the LAS (Leica Application Suite) image analysis system (Leica Microsystems, Germany). On each photographed cross-section, the number of cambium cells was counted and the widths of the developing xylem were measured along three radial directions.

The dynamics of xylem formation were analysed by fitting the Gompertz function to xylem increments (Prislan *et al*. 2018; Rathgeber, Santenoise & Cuny 2018), corrected for the previous tree ring width (Camarero, Guerrero-Campo & Gutierrez 1998; Oladi, Pourtahmasi, Eckstein & Bräuning 2011), as follows:

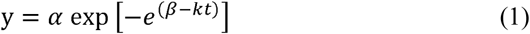

where *y* is the cumulative ring width (μm) at time *t* (day of the year), *α* is the final asymptotic size representing the annual potential growth, *β* is the x-axis placement parameter and *k* is the rate of change parameter.

For each tree and monitoring year, the following phenological xylem formation phases were recorded: *i*) cambium *reactivation*, *ii*) beginning of cell *enlargement* (bE); *iii*) beginning of cell wall *thickening* (bW); *iv*) beginning of cell *maturation* (bM); *v*) *cessation* of cell enlargement (cE); and *vi*) cessation of cell wall thickening and *lignification* (cW). The date of cambium reactivation was assessed as the average between dates when an increase of cambium cells was observed (i.e., from 3-4 to 6-7 cells in a radial row) (Čufar, Prislan, De Luis & Gričar 2008; Deslauriers, Rossi, Anfodillo & Saracino 2008). Phases of xylem growth and ring formation were computed using logistic regression, spanning from the 50% probability that phenophases have started or ended (Rathgeber *et al*. 2018). Based on phenological phases, the durations of key wood formation phases were calculated: *i*) the overall duration of the enlargement period (*d*E = cE – bE); *ii*) the duration of the wall-thickening period (*d*W = cW – bW); and *iii*) the total duration of wood formation (i.e., the duration of xylogenesis) (*d*X = cW – bE). Data were analysed using the CAVIAR (v2.10-0) package (Rathgeber *et al*. 2018) built for R statistical software (R Development Core Team 2018).

Starting from the detailed time-resolved data from tree microcores, the annual C fixed in the stem (SG_t_) was estimated for each of the sampled tree, as follows:

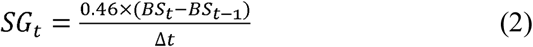

where SG_t_ is the amount of C fixed in the stem per year expressed in Mg C yr^−1^, 0.46 is the carbon content of the woody tissues (Scarascia-Mugnozza, Bauer, Persson, Matteucci & Masci 2000), BS*_t_* and BS*_t-1_* are the stem biomass in Mg of Dry Matter (DW) of at the beginning and at the end of each sampling year, Δ*t* is the time variation (one year).

The site-specific allometric equation for beech used for BS was that proposed by Masci (2002):

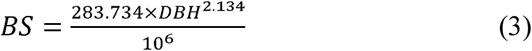

Where BS is in Mg DW, and DBH is the diameter (in cm) at 1.30 m (R^2^ = 0.96, *p*-value < 0.01).

Upscaling to stand scale was performed according to the following equation:

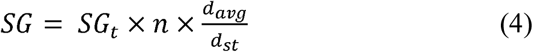

with SG expressed in Mg C ha^−1^ yr^−1^, *n* being the number of trees per hectare, d_avg_ the average diameter (cm) and d_st_ the diameter of the sampled tree (cm).

### Stem CO_2_ efflux (ES)

Two PVC collars (10 cm diameter and 5 cm high, one facing north and one south) were fixed on each tree with flexible plastic ties and sealed leak tight with Terostat (Henkel KgaA, Germany). When present, bark mosses and lichens were removed. Stem CO_2_ efflux was measured with a portable IRGA (EGM 4, PP-System, Hitchin, UK), equipped with a closed-dynamic chamber (SRC-1, PP-System, Hitchin, UK), which was tightened. Each measurement consisted of a 120-second loop, in which the CO_2_ concentration inside the chamber was measured every 5 seconds. During measurements, the CO_2_ concentration typically increased by 10 to 50 μmol mol^−1^. ES measurements were performed 16, 12, and 12 times in 2015, 2016, and 2017, respectively.

Stem CO_2_ efflux (ES) was calculated as:

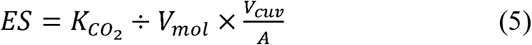

where ES is the stem CO_2_ efflux per unit surface area (μmol m^−2^ s^−1^), *K_CO2_* (μmol mol^−1^ s^−1^) is the slope of the regression between CO_2_ concentration and time during measurements, *V_mol_*, the molar volume, is the volume occupied by one mole of CO_2_ (m^3^ mol^−1^), at the air pressure (measured by built-in sensor of the EGM-4) and air temperature (T_air_ in °C) at the measurement time, *A* is the exposed lateral surface area of the stem (m^2^), and *V_cuv_* is the sum of SRC-1 and collar volumes (m^3^).

An exponential function was used to assess the relationship between ES and T_air_

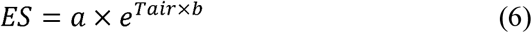

and ES overall temperature sensitivity for a 10 °C increase (Q_10_) was calculated according to Gruber *et al*. (2009) as:

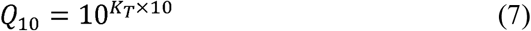

where *K_T_* is the regression slope taken from linear regression of log10 of ES versus T_air_.

From the wood formation and xylem phenology analysis described above, we identified wood (*w*) and non-wood (*nw*) formation periods for each tree, making it possible to divide the measured ES into two groups, ES_w_ and ES_nw_. According to Eq. 5, we calculated for each group the specific CO_2_ efflux at a base air temperature of 15°C (ES_15w_ and ES_15nw_) and the specific Q_10_ (Q_10w_ and Q_10nw_).

During the non-wood formation period, ES_nw_ (μmol m^−2^ s^−1^) was constituted only by the effluxes derived by maintenance respiration (ESb, μmol m^−2^ s^−1^), which was calculated as:

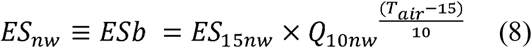

During the wood formation period, ES_nw_ (μmol m^−2^ s^−1^), which is affected by both maintenance and growth respiration, was calculated as:

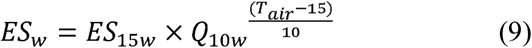

We assumed that ESb and its relationship with air temperature was also valid during the wood formation period, although this approach does not account for the acclimatisation of maintenance respiration to temperature during warmer periods (Collalti *et al*., 2018 and references therein). However, there are contrasting hypotheses on the magnitude of acclimatisation (Carey *et al*., 1997; Stockfors *&* Linder, 1998). Under the above-mentioned assumptions, we calculated the stem CO_2_ efflux due to growth respiration, ESg (μmol m^−2^ s^−1^), as:

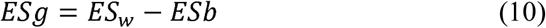

The daily C effluxes of the whole stem were obtained by integrating, over the entire stem area, the effluxes through equation 7, 8, 9, using half-hourly T_air_ values measured at the site.

The stem area was calculated as follow:

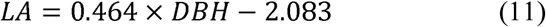

where LA is the stem lateral area (m^2^) (R^2^ = 0.828, *p*-value < 0.01, for more details on the equation see additional material Methods S1). Using DBH, we considered the measurement at 1.30 m to be representative of the whole stem, even though contrasting effects of height on stem CO_2_ effluxes have been reported (see Damesin *et al*., 2002; Katayama *et al*., 2019). Annual values of each of the C fluxes of the five sampled trees (TES, TESb, TESg, see Table 1 for definitions) were obtained by summing up the daily values.

**Table 1:**
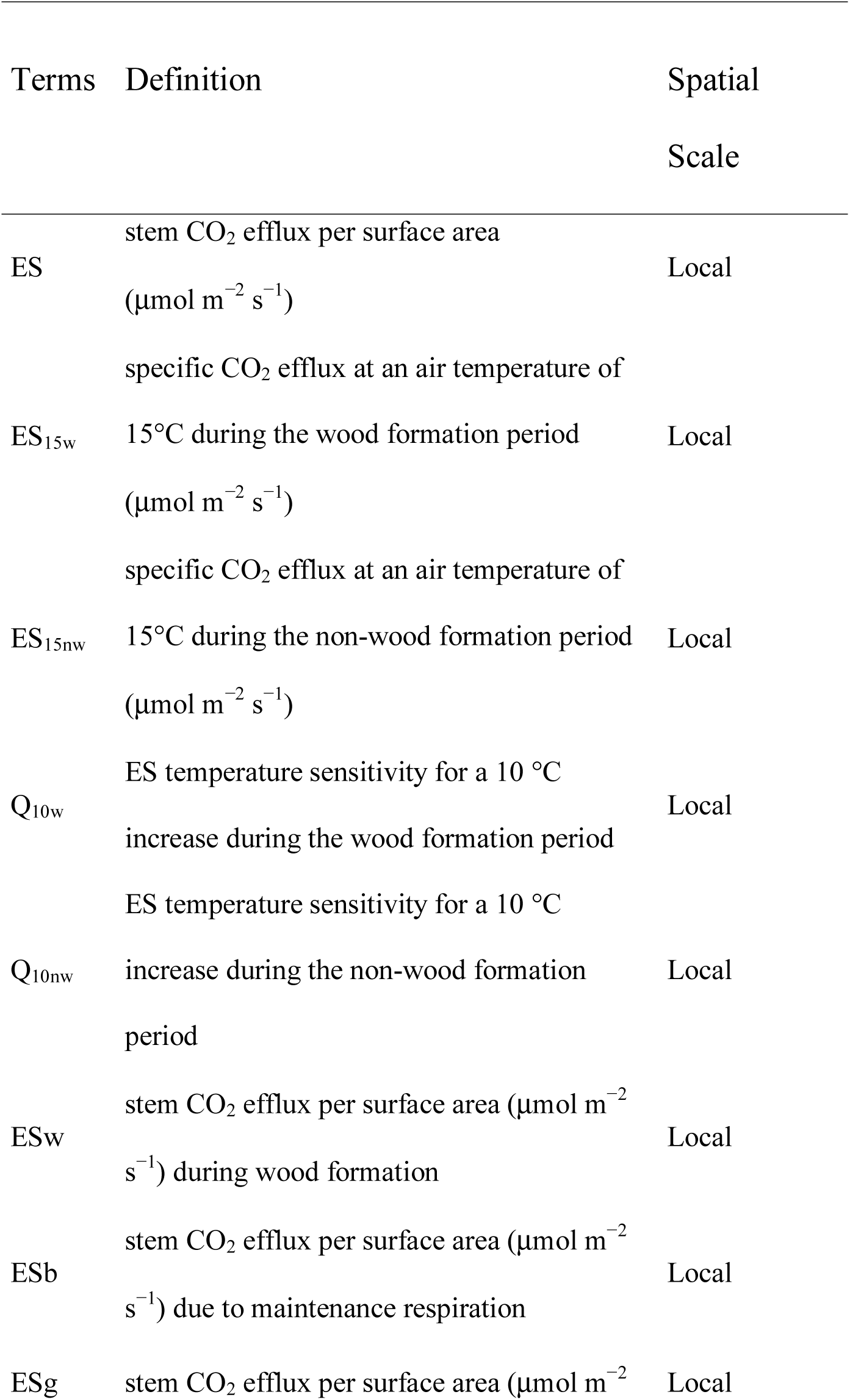
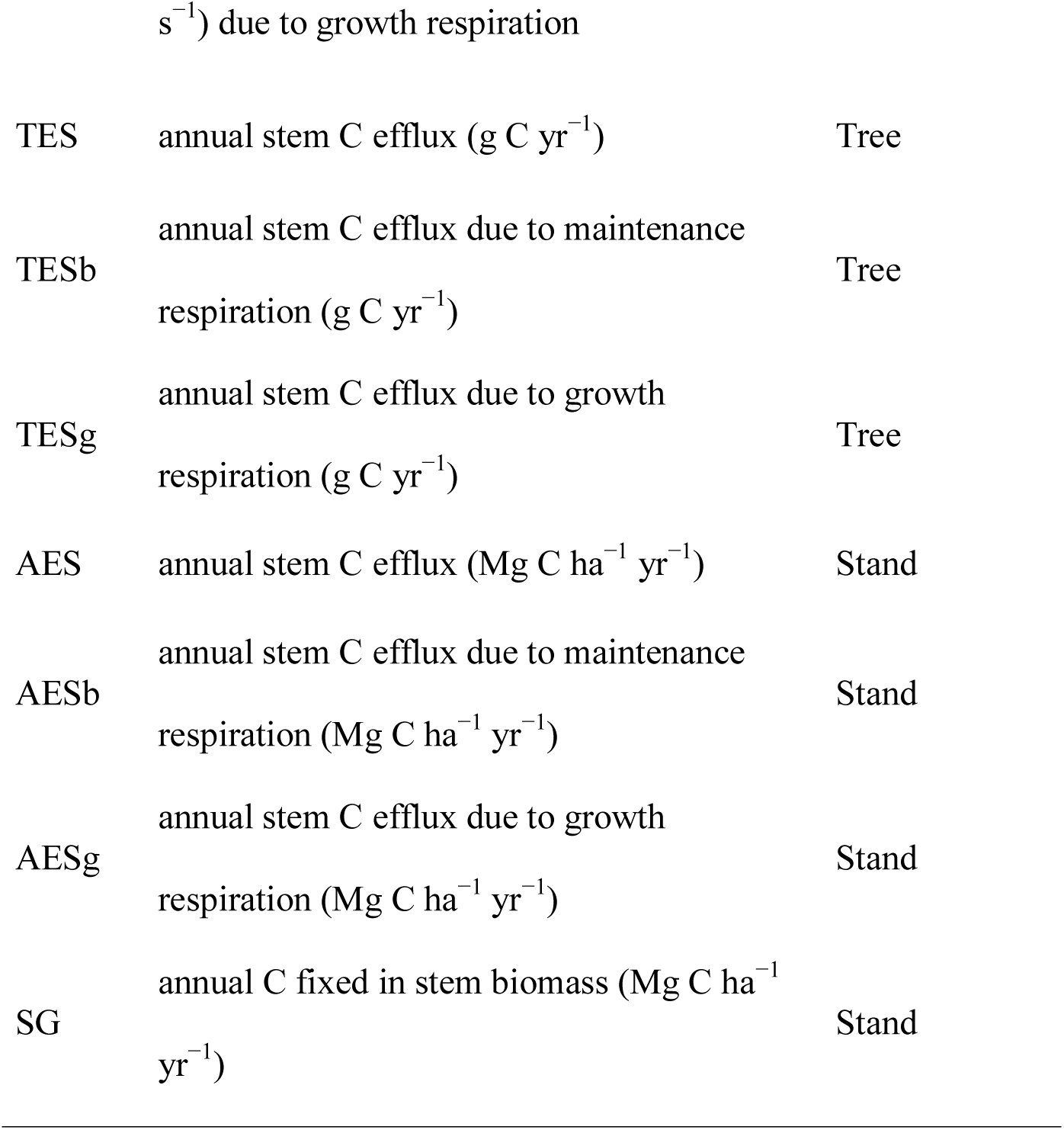
List of terms used in the text

| Terms | Definition | Spatial<br>Scale |
| --- | --- | --- |
| ES | stem CO<sub>2</sub> efflux per surface area<br>($\mu\text{mol m}^{-2} \text{s}^{-1}$) | Local |
| ES<sub>15w</sub> | specific CO<sub>2</sub> efflux at an air temperature of<br>15°C during the wood formation period<br>($\mu\text{mol m}^{-2} \text{s}^{-1}$) | Local |
| ES<sub>15nw</sub> | specific CO<sub>2</sub> efflux at an air temperature of<br>15°C during the non-wood formation period<br>($\mu\text{mol m}^{-2} \text{s}^{-1}$) | Local |
| Q<sub>10w</sub> | ES temperature sensitivity for a 10 °C<br>increase during the wood formation period | Local |
| Q<sub>10nw</sub> | ES temperature sensitivity for a 10 °C<br>increase during the non-wood formation<br>period | Local |
| ES<sub>w</sub> | stem CO<sub>2</sub> efflux per surface area ($\mu\text{mol m}^{-2}$<br>$\text{s}^{-1}$) during wood formation | Local |
| ES<sub>b</sub> | stem CO<sub>2</sub> efflux per surface area ($\mu\text{mol m}^{-2}$<br>$\text{s}^{-1}$) due to maintenance respiration | Local |
| ES<sub>g</sub> | stem CO<sub>2</sub> efflux per surface area ($\mu\text{mol m}^{-2}$ | Local |
| s<sup>-1</sup>) due to growth respiration |  |  |
| TES | annual stem C efflux (g C yr<sup>-1</sup>) | Tree |
| TESb | annual stem C efflux due to maintenance<br>respiration (g C yr<sup>-1</sup>) | Tree |
| TESg | annual stem C efflux due to growth<br>respiration (g C yr<sup>-1</sup>) | Tree |
| AES | annual stem C efflux (Mg C ha<sup>-1</sup> yr<sup>-1</sup>) | Stand |
| AESb | annual stem C efflux due to maintenance<br>respiration (Mg C ha<sup>-1</sup> yr<sup>-1</sup>) | Stand |
| AESg | annual stem C efflux due to growth<br>respiration (Mg C ha<sup>-1</sup> yr<sup>-1</sup>) | Stand |
| SG | annual C fixed in stem biomass (Mg C ha<sup>-1</sup><br>yr<sup>-1</sup>) | Stand |

**Table 2:**
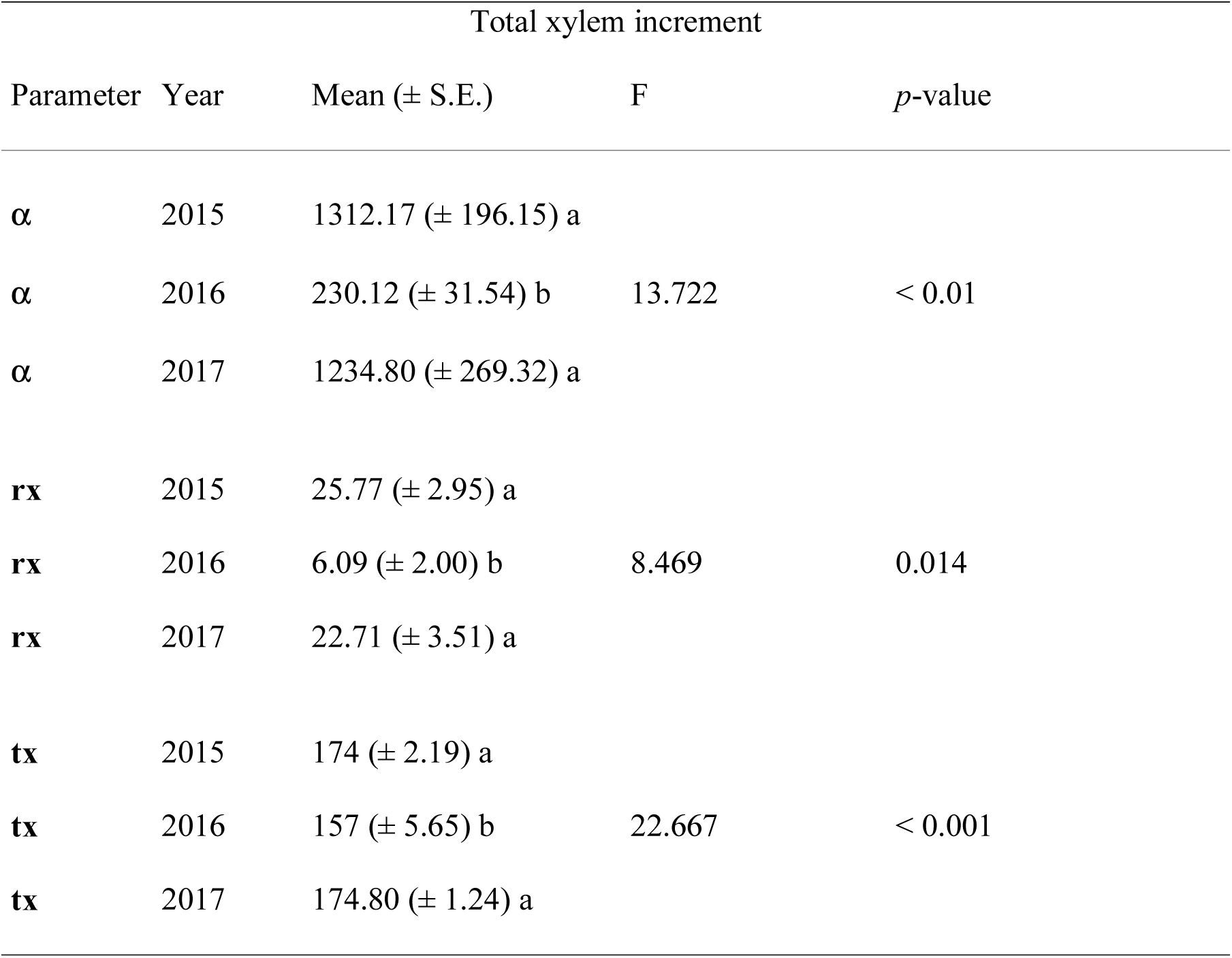
Parameters describing the intra annual radial growth derived from the Gompertz function for the total xylem increment comprised of enlarging, wall thickening and mature cells. α is the upper asymptote, representing the final ring width at the end of the growing season; tx is the DOY at which the daily increment is maximum (Gompertz curve inflection point); rx is the maximum daily increment (μm day^−1^). Different letters represent significant differences among the monitored years.

| Total xylem increment |  |  |  |  |
| --- | --- | --- | --- | --- |
| Parameter | Year | Mean (± S.E.) | F | *p*-value |
| $\alpha$ | 2015 | 1312.17 (± 196.15) a | | |
| $\alpha$ | 2016 | 230.12 (± 31.54) b | 13.722 | < 0.01 |
| $\alpha$ | 2017 | 1234.80 (± 269.32) a | | |
| **rx** | 2015 | 25.77 (± 2.95) a | | |
| **rx** | 2016 | 6.09 (± 2.00) b | 8.469 | 0.014 |
| **rx** | 2017 | 22.71 (± 3.51) a | | |
| **tx** | 2015 | 174 (± 2.19) a | | |
| **tx** | 2016 | 157 (± 5.65) b | 22.667 | < 0.001 |
| **tx** | 2017 | 174.80 (± 1.24) a | | |

On the assumption that the selected trees were representative of the stand, annual values of each of the fluxes at stand scale (AES, AESb, AESg, Mg C ha^−1^ yr^−1^, see Table 1 for definitions) were calculated:

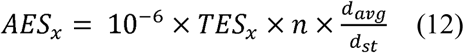

where TES_x_ was the effluxes at tree level (TES, TESb, TESg, g C yr^−1^), *n* is the number of trees per hectare, d_avg_ is the average diameter (cm) and d_st_ is the diameter of the sampled tree (cm).

### Meteorological and phenological data

For the period 1989-2014, FLUXNET2015 release half-hourly air temperature and precipitation were used (Reyer *et al*. 2019). For the period of the study (2015-2017), measured data were gap filled using downloaded data by the ERA5 database of the European Centre for Medium-Range Weather Forecasts (ECMWF) (https://www.ecmwf.int/en/forecasts/datasets/archive-datasets/reanalysis-datasets/era5), according to FLUXNET 2015 release equations.

The Standardized Precipitation Evapotranspiration Index (SPEI), considered the most appropriate index for the Mediterranean climate (Vicente-Serrano *et al*. 2013), was used to assess the magnitude of the drought in 2017. This index is based on the difference between precipitation and potential evapotranspiration (PET), computed according to Hargreaves’ equation. The 3-month SPEI was calculated for the site for the period 1989-2017, using the SPEI package in R.

Leaf phenology was monitored using the MODIS Leaf Area Index product (LAI, MOD15A2H, https://modis.gsfc.nasa.gov/) with 8-day temporal resolution and 500-meter spatial resolution. The date of onset of photosynthetic activity (green up) and the date at which plant green leaf area peaked at its annual maximum (maturity) were assessed through the rate of change in the curvature of the fitted logistic models (Zhang *et al*. 2003).

### Statistical data analysis

Descriptive parameters of growth and xylem phenology were tested using one-way repeated measures analysis of variance, with years as factor, followed by post-hoc (Holm-Sidak method). An exponential equation was used to evaluate the relationship between ES and T_air_. Differences among ES parameters (Q_10_ and ES_15_) were tested using two-way repeated measures Anova (two factor repetition), using year and period (non-wood formation, wood formation) as predictive factors. Multiple comparisons were performed by the Holm-Sidak method. Linear regressions were used to assess the relationship between tree ring widths (TRW) and mean annual ES. We tested data normality and constant variance using the Shapiro-Wilk test and the Spearman rank correlation between the absolute values of the residuals and the observed value of the dependent variable.

## Results

### Wood formation dynamics

The date of onset of photosynthetic activity, based on leaf area index (LAI) dynamics, differed among the study years, occurring at DOY 118, 95 and 127 in 2015, 2016 and 2017, respectively. In all three years, cambium reactivation occurred after leaf unfolding at DOY 123 ± 4, 118 ± 8, 138 ± 6 in 2015, 2016 and 2017, respectively (Fig. 2). In 2016, cambium cell production also continued after the late frost event, but at considerably lower rates.

**Fig. 2:**
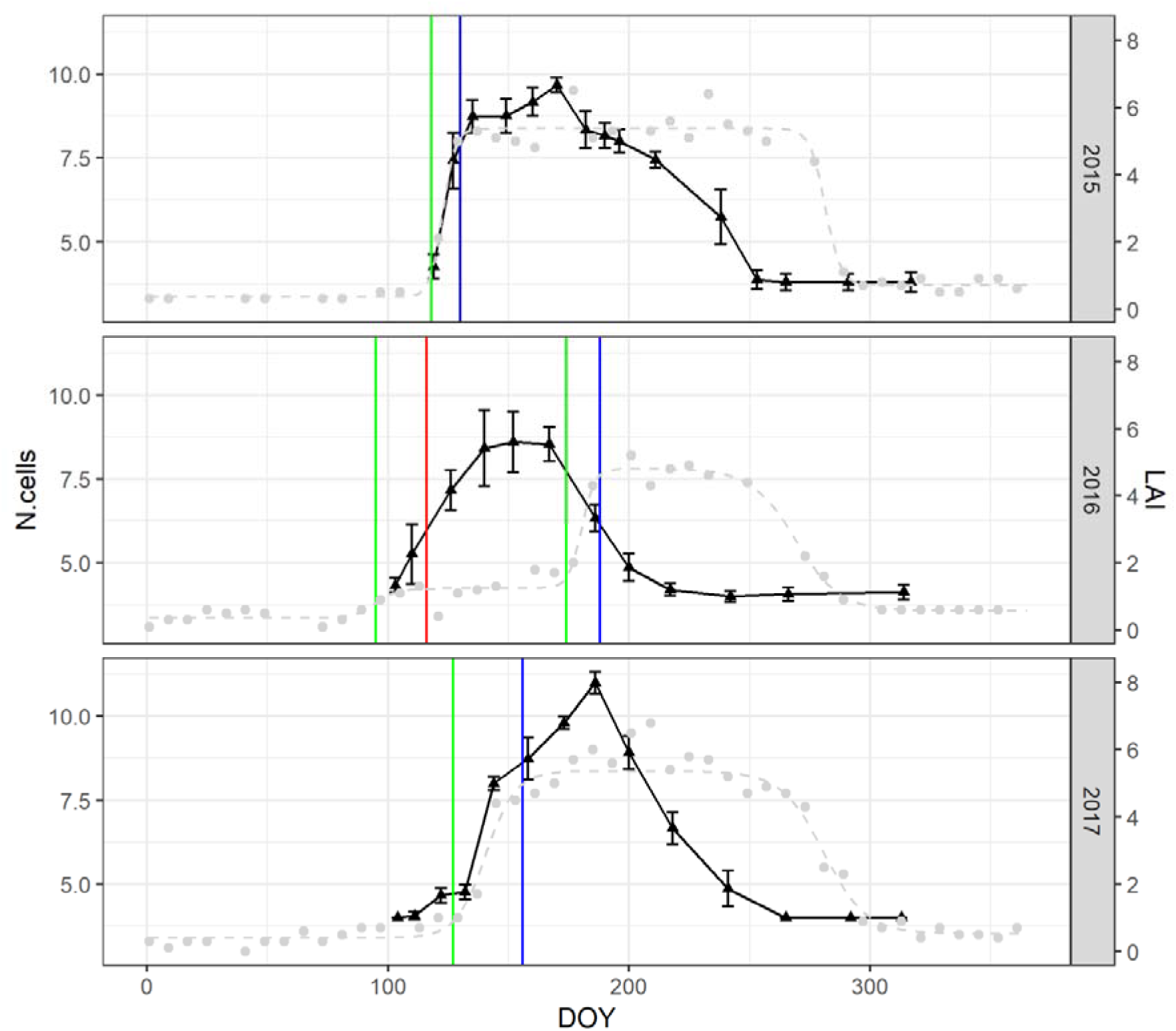
Number of cambium cells (N. cells vs. leaf area index (LAI, m^2^m^−2^)). Grey points and dashed lines are the MODIS-LAI values and modelled intra-annual dynamic of Selva Piana beech forest, respectively. Green and blue vertical lines represent the green-up and maturity phases of leaf phenology, respectively. The red vertical line represents the late frost of 25^th^ April 2016. Black triangles are the average number of cambial cells of five beech trees. Bars are the standard error.

Different intra-annual growth patterns were observed during the three study years, especially in the year of the late frost (2016, Fig. 3, Table 1). In 2016, the maximum growth rate (rx) (F = 8.469, *p*-value = 0.014) was lower and was reached 3 weeks earlier (tx) (F = 22.667, *p*-value < 0.001) than in the other two years. The different intra-annual growth patterns also resulted in significantly narrower tree rings in 2016 (230.12 ± 1.54 μm) (F = 13.272, *p*-value < 0.01) than in 2015 (1312.17 ± 196.15 μm) and 2017 (1234.80 ± 269.32 μm).

**Fig. 3:**
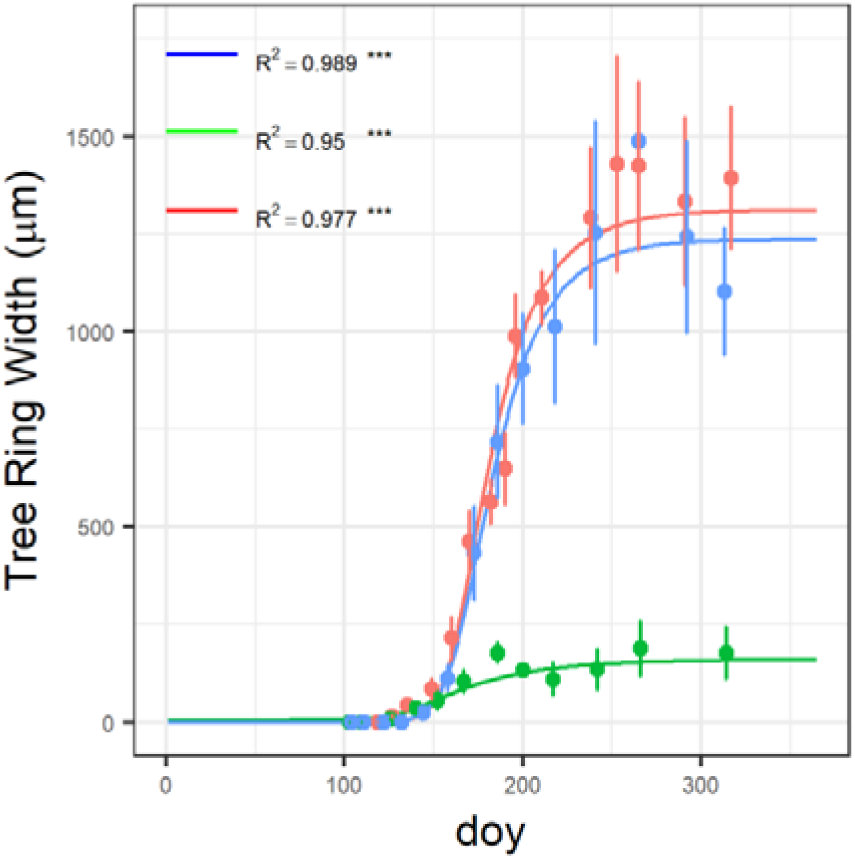
Intra-annual dynamics of xylem formation (μm) in 2015 (red dots and solid line), 2016 (green dots and solid line) and 2017 (blue dots and solid line). Gompertz functions were fitted to the total xylem increment comprised of enlarging, wall thickening and mature cells. Each point is the mean of 5 sampled *Fagus sylvatica* trees and bars are standard errors. *** *p*-value < 0.001

Differences were also observed for the beginning, cessation, and duration of wood formation phases (Fig. S2). The beginning of the enlargement phase occurred earliest in 2016 and latest in 2017 (F = 34.789, *p*-value < 0.001). In contrast, the cessation of this phase was observed latest in 2015 (F = 17.155, *p*-value < 0.01). Consequently, the duration of the enlargement phase was longer in 2015 (110 ± 22 days) than in 2016 (82 ± 4 days) and 2017 (78 ± 4 days) (F = 8.025, *p*-value = 0.01).

The beginning of the wall thickening phase did not differ among the years (F = 4.188, *p*-value = 0.06). The cessation of this phase occurred latest in 2015 (F = 69.167, *p*-value < 0.001). The duration of the wall thickening phase was thus shorter in 2016 (57 ± 5 days) than in 2015 (99 ± 6 days) and 2017 (75 ± 4 days) (F = 26.561, *p*-value < 0.001). We also observed a delay in the beginning of cell maturation in 2016 (at DOY 200 ± 4) with respect to 2015 (at DOY 178 ± 4) and 2017 (at DOY 176 ± 2) (F = 11.650, *p*-value < 0.01). The overall duration of wood formation was longer in 2015 (128 ± 5 days) than in 2016 (98 ± 8 days) and 2017 (97 ± 8 days) (F = 12.561, *p*-value < 0.001).

### Stem CO_2_ efflux (ES)

During the monitoring period (April 2015 – November 2017), the measured ES ranged between 0.16 ± 0.03 μmol CO_2_ m^−2^ s^−1^ (December 2015) and 3.01 ± 0.40 μmol CO_2_ m^−2^ s^−1^ (August 2017) (Fig. 5). Mean ES measured in 2016 (0.68 ± 0.19 μmol CO_2_ m^−2^ s^−1^) was lower (F = 24.476, *p*-value < 0.01) than in 2015 (1.11 ± 0.40 μmol CO_2_ m^−2^ s^−1^) and 2017 (1.29 ± 0.30 μmol CO_2_ m^−2^ s^−1^).

**Fig. 4:**
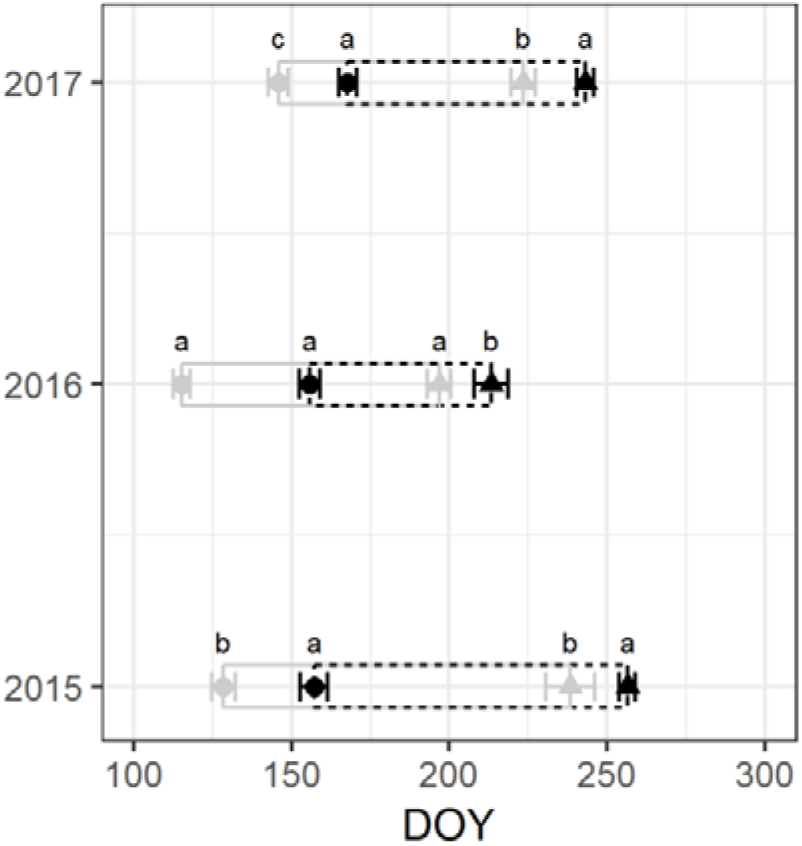
Critical dates and duration of wood formation phases. Different letters (a, b and c) represent significant differences among the beginning of the enlargement phase (grey dots), the beginning of the wall thickening phase (black dots), cessation of the enlargement phase (grey triangle) and cessation of the wall thickening phase (black triangle) (*p-*value < 0.05). Grey and black rectangles represent the duration of the enlargement and wall thickening phases, respectively. The sum of their length is the duration of wood formation. Each shape is the mean of 5 sampled trees per year and the bars error represent standard error.

**Fig. 5:**
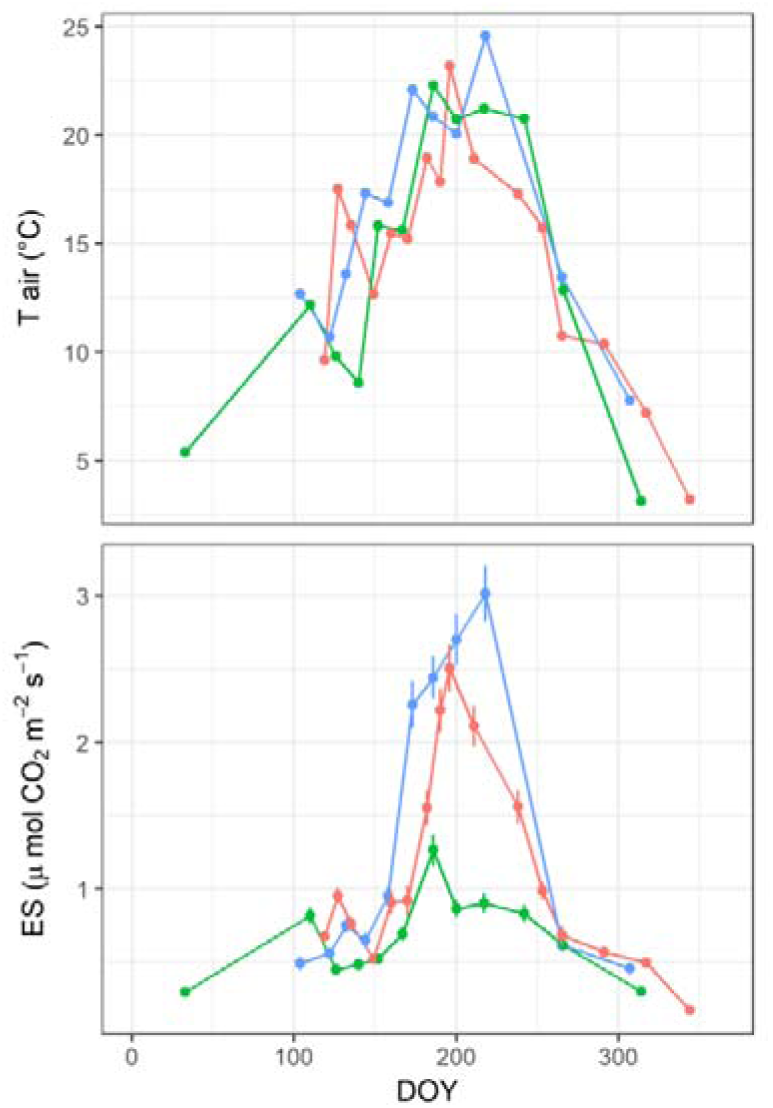
Top panel: T_air_ (°C) at the measuring time in 2015 (red), 2016 (green) and 2017 (blue). Bottom panel: Measured stem CO_2_ effluxes (μmol CO_2_ m s) in 2015 (red), 2016 (green) and 2017 (blue). Each point is the mean of 5 *Fagus sylvatica* trees. Bars are standard errors.

In each year, ES was strongly related to air temperature through the standard exponential function (Fig. 6). The relation was confirmed at different aggregation levels of measurements (whole year, wood formation and non-wood formation periods; see also Table S1 in Supporting Information).

**Fig. 6:**
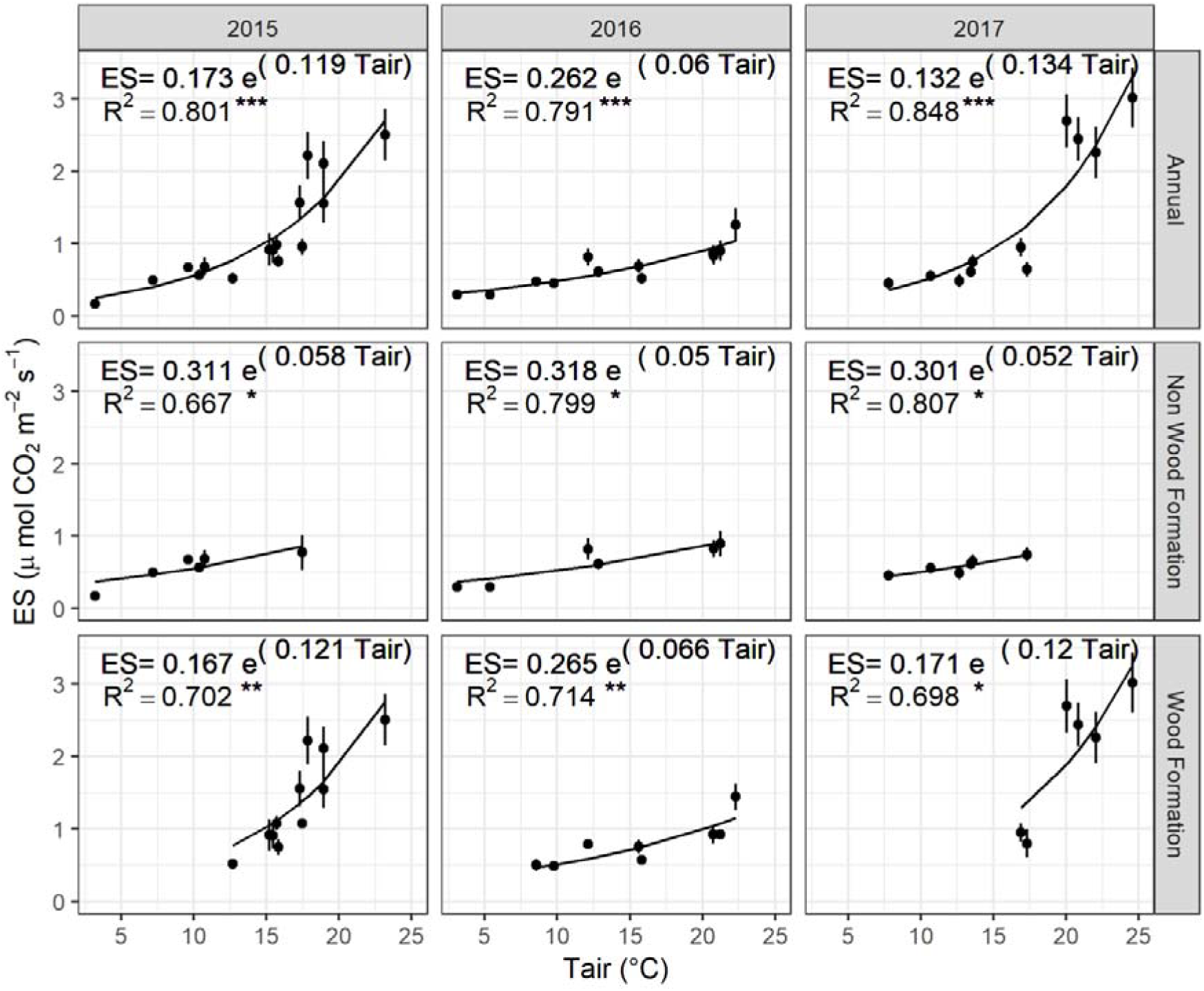
Relationship between ES (μmol CO_2_ m s) and air temperature (T_air_, °C). Annual, considering the whole measurements for each year, each point is the mean of five sampled trees. Each point represents the mean of those trees, during non-wood and wood formation phases, at a given sampling date. Bars are the standard error. *** *p*-value < 0.001, ** *p*-value < 0.01, * *p*-value < 0.05.

Average Q_10_ was 2.71 ± 0.15, 2.11 ± 0.18 and 2.68 ± 0.15, in 2015, 2016 and 2017, respectively. The Q_10_ parameter was not strongly affected by the sampling year (*p*-value = 0.059), although the values in 2016 were 22% lower than in the other two years. Wood formation affected the Q_10_ parameter (F = 31.563, *p*-value < 0.01) with Q_10w_ and Q_10nw_ calculated to be 3.06 ± 0.15 and 1.93 ± 0.14 (t = 5.571, *p*-value < 0.01), respectively. This difference was confirmed for all of the sampled years.

ES_15_ was also affected by the different conditions of the monitoring years (F = 7.094, *p*-value = 0.01) with mean values in 2016 (0.63 ± 0.07 μmol CO_2_ m^−2^ s^−1^) lower than in 2015 (0.93 ± 0.12 μmol CO_2_ m^−2^ s^−1^) and 2017 (0.82 ± 0.12 μmol CO_2_ m^−2^ s^−1^). Similar to Q_10_, the wood formation period also affected ES_15_, with ES_15w_ (0.84 ± 0.22 μmol CO_2_ m^−2^ s^−1^) higher than ES_15nw_ (0.73 ± 0.02 μmol CO_2_ m^−2^ s^−1^, F = 7.094, *p*-value = 0.01). Furthermore, during the wood formation period, ES_15w_ was higher in 2015 (1.03 ± 0.07 μmol CO_2_ m^−2^ s^−1^) and 2017 (0.90 ± 0.07 μmol CO_2_ m^−2^ s^−1^) than in 2016 (0.60 ± 0.09 μmol CO_2_ m^−2^ s^−1^). No differences among years were found for ES during the non-wood formation periods.

Annual ES for individual trees, per unit of lateral surface area, ranged between 112 g C m^−2^ yr^−1^ in 2016 (tree 4) and 349 g C m^−2^ yr^−1^ in 2017 (tree 2). Average total ES for all sampled trees was lower in 2016 (182 ± 25 g C m^−2^ yr^−1^, F = 12.007, *p*-value < 0.01) than in 2015 (258 ± 27 g C m^−2^ yr^−1^) and 2017 (233 ± 35 g C m^−2^ yr^−1^).

The estimated contribution of maintenance respiration to ES for individual trees ranged between 112 g C m^−2^ yr^−1^ in 2016 (tree number 4 showed no growth) and 284 g C m^−2^ yr^−1^ in 2017 (tree number 2), and was lower, on average, in 2016 (169 ± 21 g C m^−2^ yr^−1^) than in 2015 (211 ± 18 g C m^−2^ yr^−1^) (q = 5.104, *p*-value = 0.017).

Likewise, the estimated contribution of wood formation to ES for individual trees varied between 0 in 2016 (tree number 4) and 70 g C m^−2^ yr^−1^ in 2015 (tree number 2) and was significantly lower (F = 8.144, *p*-value = 0.012) on average in 2016 (14 ± 5 g C m^−2^ y^−1^) than in 2015 (48 ± 9 g C m^−2^ yr^−1^) and 2017 (39 ± 8 g C m^−2^ yr^−1^). In relative terms, the contribution of wood formation to ES was estimated to be 18 ± 2%, 9 ± 3% and 16 ± 3% in 2015, 2016 and 2017, respectively, with the remaining CO_2_ efflux originating from maintenance respiration.

### Radial growth and stem C effluxes

During the study period, annual average measured ES and tree ring widths were closely related (Fig. 5). Seasonal patterns of ES were similar in the three years, but with different amplitudes (Fig. 8). Moreover, ESb, the stem C effluxes affected by maintenance respiration, showed a similar pattern among the three years. We observed a time-lag between the time of maximum growth rate (tx) and maximum ES values of 23 ± 2 days, 31 ± 2 days and 29 ± 1 days in 2015, 2016 and 2017, respectively. Differences between years were not significant (F = 3.317, *p*-value = 0.07).

**Fig. 7:**
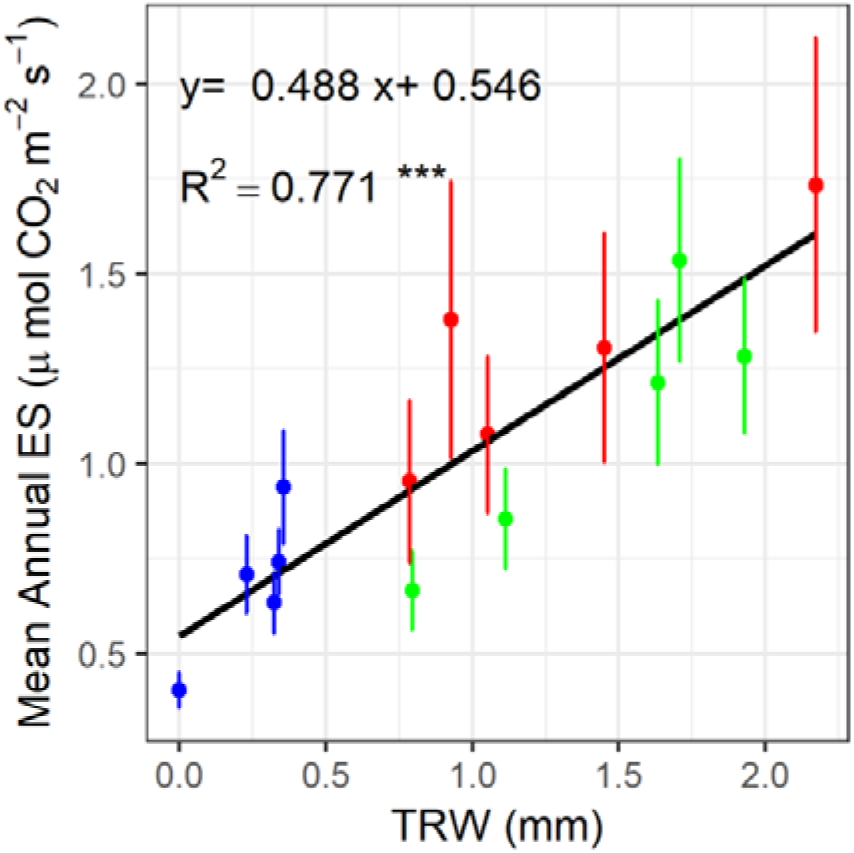
Relationships between ring widths (mm) and the mean annual ES (μmol CO_2_ m s) measured in 2015 (green), 2016 (blue), and 2017 (red). Each point represents a sampled *Fagus sylvatica* tree per year. Bars are the standard error. *** *p*-value < 0.001.

**Fig. 8:**
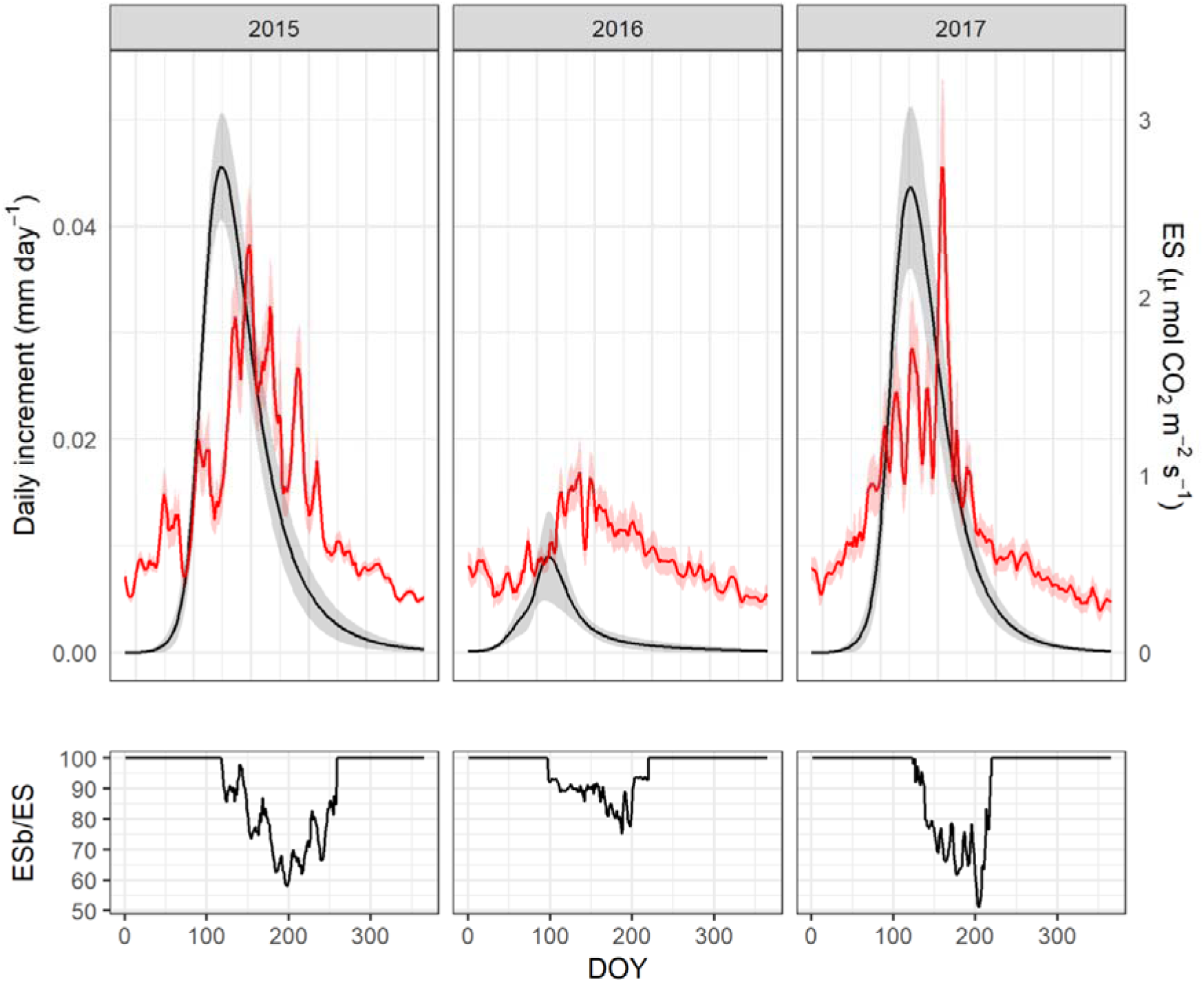
Daily increment and stem Carbon effluxes of *Fagus sylvatica*. The black line is the daily increment during 2015, 2016 and 2017, respectively. The red line represents ES, the daily C effluxes. The lower panel, ESb/ES shows the contribution of maintenance respiration to daily stem C efflux.

### From tree to stand level

Annual stand-level stem C emissions (AES) were lower in 2016 (1.58 ± 0.22 Mg C ha^−1^ yr^−1^) than in 2015 (2.24 ± 0.24 Mg C ha^−1^ yr^−1^) and 2017 (2.02 ± 0.27 Mg C ha^−1^ yr^−1^) (Table 3). Annual stem C effluxes due to maintenance respiration (AESb) in 2015 (1.47 ± 0.19 Mg C ha^−1^ yr^−1^) were lower than in 2016 (1.82 ± 0.16 Mg C ha^−1^ yr^−1^); AES due to growth respiration (AESg) was lower in 2016 (0.11 ± 0.04 Mg C ha^−1^ yr^−1^) than in 2015 (0.42 ± 0.08 Mg C ha^−1^ yr^−1^) and 2017 (0.34 ± 0.06 Mg C ha^−1^ yr^−1^). The contribution of AESg to the annual stem effluxes was 19%, 7% and 16% in 2015, 2016 and 2017, respectively.

**Table 3:** Annual C stem fluxes. AES is the annual stem C efflux assessed using specific parameters for wood formation (Q_10w_ and ES_15w_) and non-wood formation phases (Q_10nw_ and ES_15nw_); AESb is the annual stem C efflux due to maintenance respiration; AESg is the annual stem C efflux due to growth respiration; SG is the annual amount of C fixed in the stem biomass; different letters represent significant differences *p*-value < 0.05

| Year | Flux type | Mean ± (S.E.) Mg C ha<sup>-1</sup> y<sup>-1</sup> | F | $p$-value |
| --- | --- | --- | --- | --- |
| 2015 | AES | 2.24±(0.24) a | 11.800 | 0.004 |
| 2016 | AES | 1.58±(0.22) b | | |
| 2017 | AES | 2.02±(0.27) a | | |
| 2015 | AESb | 1.82±(0.16) a | 8.145 | 0.012 |
| 2016 | AESb | 1.47±(0.19) b | | |
| 2017 | AESb | 1.73±(0.18) a/b | | |
| 2015 | AESg | 0.42±(0.08) a | 7.573 | 0.014 |
| 2016 | AESg | 0.11±(0.04) b | | |
| 2017 | AESg | 0.34±(0.06) a | | |
| 2015 | SG | 2.78±(0.41) a | 16.534 | 0.001 |
| 2016 | SG | 0.48±(0.13) b | | |
| 2017 | SG | 2.50±(0.49) a | | |

The amount of carbon fixed in the stem biomass (SG) was lower in 2016 (0.48 ± 0.13 Mg C ha^−1^ yr^−1^) than in 2015 (2.78 ± 0.41 Mg C ha^−1^ y^−1^) and 2017 (2.50± 0.49 Mg C ha^−1^ yr^−1^). At the studied beech forest, the mean C construction cost of wood, defined as the slope of the relationship between AESg and SG at tree level (R^2^ = 0.849, *p*-value < 0.01, see Supporting Information Fig.S3), was 0.2 g C emitted per g C fixed. On an annual scale, this parameter was 0.15 ± 0.01 for 2015, 0.24 ± 0.05 for 2016 and 0.14 ± 0.02 for 2017.

## Discussion

### Cambial activity and radial growth are not entirely linked to leaf phenology

To the best of our knowledge, this is the first time that the effects of a spring late frost with a subsequently close summer drought on the wood formation dynamic of beech have been described.

In all three years, cambium reactivation and wood formation occurred within 1-3 weeks after leaf development, confirming the tight dependence of radial growth on leaf phenology and photosynthesis in diffuse-porous species (Čufar *et al*. 2008; Michelot *et al*. 2012). Nevertheless, in diffuse-porous trees, stem conductivity to water occurs in several outermost growth rings and is not limited to the youngest formed xylem, as found in ring-porous species (Schume, Grabner & Eckmüllner 2004). Hence, in beech allocation to the current year wood, it is not as decisive as in ring-porous species, and newly formed photosynthates at the beginning of the season are preferably used for other crucial processes, such as foliage and fine root growth. At the same site, Matteucci (1998) analysed in parallel net ecosystem exchange (NEE) and carbon allocation to foliage and stem radial growth, finding that the latter started approximately 15 days after photosynthesis exceeded respiration (i.e., NEE was negative). Until then, net absorbed carbon was allocated mostly to foliage growth. This can be related to the C allocation hierarchy, which identifies newly developing leaves as the main C sink at the beginning of the growing season (Campioli *et al*., 2013; Collalti *et al*., 2016; Merganičová *et al*., 2019) Interestingly, and even counter to our expectations, in 2016, the cambium remained active at low rates even after complete canopy defoliation, probably fuelled by old C reserves (D’Andrea *et al*. 2019).

After the second re-sprouting, cambium cell production decreased and became non-productive, although the environmental conditions were potentially still favourable for radial growth, being related to day-length (Rossi *et al*. 2006; Camarero, Olano & Parras 2010). Stem radial growth of beech in Selva Piana was greatly affected by the extreme late spring frost in 2016 because of the premature cessation of cambial cell production and the lower growth rate during the active period, which resulted in 82% narrower annual xylem increments compared to 2015 and 2017. This can be related to a somewhat hypothesised, genetically controlled, form of hierarchy in C allocation (composed by old C reserve and recently fixed photosynthates), which identifies newly developing leaves as the main C sink rather than radial growth. In beech, previously reported growth reduction – as a consequence of late frost – has ranged from 48 to 83% in beech, with the maximum occurring at the northern fringe of the Alps (Dittmar *et al*. 2006). Radial growth rates had fully recovered in 2017, with no visible long-term effects of the late spring frost event in 2016, showing the high resilience of beech growth to late frost (Dittmar *et al*. 2006; Principe, Struwe, Wilmking & Kreyling 2017). However, as recently shown in D’Andrea *et al*. (2019), beech trees during 2016 were able to compensate the lost C reserve, completely refilling the pool to the same level as before the frost event. Hence, there was no need to prioritize reserve recharge over stem biomass production the subsequent year.

The 2017 summer drought became severe only in August (SPEI > 1.5), when the trees had already completed most of their radial growth, as already seen for other tree species growing in the Mediterranean adjusting the end xylem growth before potential stressful conditions may occur (e.g. Lempereur *et al*., 2015; Forner *et al*., 2018). Instead, the importance of spring climatic conditions on beech growth has been reported for the Apennines and the Eastern Alps (Piovesan, Biondi, Di Filippo, Alessandrini & Maugeri 2008; Di Filippo, Biondi, Maugeri, Schirone & Piovesan 2012).

### Effluxes from stem are not entirely synchronised to radial growth

Mean annual values of Q_10_ ranged between 2.11 (2016) and 2.71 (2015) and were similar to the values estimated at the same site for co-dominant (2.59) and dominant (2.34) trees (Guidolotti *et al*. 2013). Q_10nw_ and Q_10w_ estimated in this study are closely comparable with the dataset of various coniferous and broadleaf tree species reported in Damesin et al. (2002). Similar intra-annual variability of Q_10_ has been observed in many studies on different species with higher Q_10_ during the growing period (Paembonan, Hagihara & Hozumi 1992; Carey, Callaway & DeLucia 1997; Stockfors & Linder 1998; Gruber *et al*. 2009). However, other studies found (or assume) stable Q_10_ throughout the year (Ceschia *et al*. 2002; Damesin *et al*. 2002).

Our results suggests that wood formation affects Q_10_, indeed Q_10w_ was higher than Q_10nw_, despite higher temperatures during the wood formation period, exactly contrary to a case of sole control of temperature on the parameters (Atkin & Tjoelker 2003).

As reported in other studies, ES15, stem CO_2_ efflux at an air temperature of 15°C, was sensitive to wood formation processes, showing an increase during the growing period (Ceschia *et al*. 2002; Damesin *et al*. 2002).

Maximum xylem production and maximum ES were not synchronized, while a constant delay, of about a month, was observed, as similarly found in a young beech forest in which peak ES occurred *c*. 27 days after the maximum stem growth rate (Ceschia *et al*. 2002). Furthermore, our results confirmed that peak ES occurred when xylem cells were still in the phase of wall thickening and lignification, as previously hypothesized (Ceschia *et al*. 2002). Moreover, when maximum ES was observed, it is very likely that trees were already refilling the stem C reserves pool (Scartazza, Moscatello, Matteucci, Battistelli & Brugnoli 2013).

### Only spring frost affects negatively stem C fluxes

The amount of C fixed by stem biomass formation in 2015 and 2017 ranged from 2.58 to 2.70 Mg C ha^−1^ yr^−1^, a bit higher than the values reported for a beech forest in Germany, ranging from 1.69 to 2.41 Mg C ha^−1^ yr^−1^ (Mund *et al*. 2010). In 2016, however, we measured only 0.48 ± 0.13 Mg C ha^−1^ yr^−1^, i.e., only about 20% of the fixation during the two reference years, emphasizing how exceptionally negative this year was.

Annual stem CO_2_ efflux (AES) is known to be highly variable in temperate forests (Yang *et al*. 2016). Our data ranges from 1.58 to 2.24 Mg C ha^-1^ yr^-1^ and thus is similar to the 1.65 to 2.25 Mg C ha^−1^ yr^−1^ reported for a younger beech forest (Damesin *et al*. 2002).

An earlier estimate of AES at the study site was 0.63 Mg C ha^-1^ yr^−1^ (Guidolotti *et al*. 2013) for 2007, year characterized by an extreme summer drought (SPEI > 2). The stem C efflux of the drought year presented in this study (2017, 1.58 Mg C ha^−1^ yr^−1^) was about 150% higher than in 2007, which could be due to an increase in stem biomass (*c*. 15% lower in 2007 than in 2017, see also Collalti *et al*., 2019) and to different measurement tools. The contribution of AESg to annual stem effluxes, ranging from 7% to 19%, was lower than that measured in a young beech forest (Ceschia et al., 2002), evidencing the importance of forest developmental stage in determining wood formation and growth respiration. The construction cost we found of 0.23 g C fixed per g C emitted is consistent with the values reported for beech, 0.2 – 0.38 g C fixed per g C emitted (Ceschia *et al*. 2002; Damesin *et al*. 2002).

While the late frost event in 2016 reduced both wood growth and stem CO_2_ efflux with respect to those measured in the other two years, the percentage of growth reduction (80%) was much larger than the reduction of ES (25%). Hence, it seems that in mature beech the contribution of growth respiration to total stem CO_2_ fluxes is lower than that of maintenance respiration, as reported in Collalti *et al*.(2019). In the year of late frost, the strong reduction of fixed growth C and the contemporary lower reduction of stem CO_2_ efflux strongly affected the overall stem carbon balance. In contrast, the summer drought did not have any effect on stem growth, and thus neither on C efflux due to growth respiration.

In conclusion, this study further highlights the sensitivity of beech to leaf damage due to late spring frost. Since leaf development is forecast to start earlier for beeches due to global warming (Augspurger 2013), the likelihood that spring frost may damage leaves will increase. We demonstrated that stem growth was significantly reduced due to the prolonged absence of photosynthesizing leaves after frost, since beech trees tapped their pool of old C reserves. However, the loss in growth was not completely compensated for after re-growth of leaves, but rather the cambium activity ceased shortly thereafter. Consequently, the trees fixed less C in the stem biomass, showing also a reduction of the stem carbon efflux related to growth respiration.

Moreover, the summer drought occurred too late to affect wood formation and stem CO_2_ effluxes. However, more investigations are needed to evaluate its effects on other physiological processes. This study also underlines the crucial role of spring weather conditions on the growth and physiology of beech trees.

A better understanding of fine scale C dynamic will help in evaluating a medium- to long-term response to climate change under an increasing frequency of extreme events.

## Acknowledgements

The activities of Negar Rezaie at the wood anatomy laboratory of Slovenian Forestry Institute were supported by an Excellence Research Award of the National Research Council of Italy, Department of Biology, Agriculture, and Food Secures (Prot. 71951, 06/11/2017). Collelongo-Selva Piana is one of the sites of the Italian Long Term Ecological Research network (LTER-Italy), part of the International LTER network (ILTER). Research at the site in the years of this study was funded by the eLTER H2020 project (grant agreement no. 654359). Authors are grateful to Martin for English language editing.

## Author contribution

E.D’A., N.R., G.M. contributed to the design of the research. Fieldwork was carried out by E.D’A., N.R.; laboratory analysis for wood formation N.R., J.M., P.P.; data analysis was done by E.D’A.; data interpretation by E.D’A., N.R., P.P., J.M., A.C., G.M., J.K.. The manuscript was written by E.D’A and N.R. with major contributions by P.P., J.M., A.C., G.M., J.K..

## References

Allevato E., Saulino L., Cesarano G., Chirico G.B., D’Urso G., Falanga Bolognesi S., … Bonanomi G. (2019) Canopy damage by spring frost in European beech along the Apennines: effect of latitude, altitude and aspect. Remote Sensing of Environment 225, 431–440.

Atkin O.K. & Tjoelker M.G. (2003) Thermal acclimation and the dynamic response of plant respiration to temperature. Trends in Plant Science 8, 343–351.

Augspurger C.K. (2013) Reconstructing patterns of temperature, phenology, and frost damage over 124 years□: Spring damage risk is increasing. Ecology 94, 41–50.

Bascietto M., Bajocco S., Mazzenga F. & Matteucci G. (2018) Assessing spring frost e ff ects on beech forests in Central Apennines from remotely-sensed data. Agricultural and Forest Meteorology 248, 240–250.

Bascietto M., Cherubini P. & Scarascia-Mugnozza G. (2004) Tree rings from a European beech forest chronosequence are useful for detecting growth trends and carbon sequestration. Canadian Journal of Forest Research 34, 481–492.

Begum S., Nakaba S., Oribe Y., Kubo T. & Funada R. (2007) Induction of Cambial Reactivation by Localized Heating in a Deciduous Hardwood Hybrid Poplar (Populus sieboldii 3 P. grandidentata). Annals of botany, 439–447.

Bloemen J., Agneessens L., Van Meulebroek L., Aubrey D.P., Mcguire M.A., Teskey R.O. & Steppe K. (2014) Stem girdling affects the quantity of CO2 transported in xylem as well as CO2 efflux from soil. New Phytologist 201, 897–907.

Bloemen J., McGuire M.A., Aubrey D.P., Teskey R.O. & Steppe K. (2013) Transport of root-respired CO 2 via the transpiration stream affects aboveground carbon assimilation and CO 2 efflux in trees. New Phytologist 197, 555–565.

Bowman W.P., Barbour M.M., Turnbull M.H., Tissue D.T., Whitehead D. & Griffin K.L. (2005) Sap flow rates and sapwood density are critical factors in within- and between-tree variation in CO2 efflux from stems of mature Dacrydium cupressinum trees. New Phytologist 167, 815–828.

Camarero J.J., Guerrero-Campo J. & Gutierrez E. (1998) Tree-Ring Growth and Structure of Pinus uncinata and Pinus sylvestris in the Central Spanish Pyrenees. Arctic and Alpine Research 30, 1.

Camarero J.J., Olano J.M. & Parras A. (2010) Plastic bimodal xylogenesis in conifers from continental Mediterranean climates. New Phytologist 185, 471–480.

Campioli M., Verbeeck H., Van den Bossche J., Wu J., Ibrom A., D’Andrea E., … Granier A. (2013) Can decision rules simulate carbon allocation for years with contrasting and extreme weather conditions? A case study for three temperate beech forests. Ecological Modelling 263.

Carey E. V., Callaway R.M. & DeLucia E.H. (1997) Stem respiration of ponderosa pines grown in contrasting climates: implications for global climate change. Oecologia 111, 19–25.

Carrer M., Brunetti M. & Castagneri D. (2016) The Imprint of Extreme Climate Events in Century-Long Time Series of Wood Anatomical Traits in High-Elevation Conifers. Frontiers in Plant Science 7, 1–12.

Ceschia É., Damesin C., Lebaube S., Pontailler J.Y. & Dufrêne É. (2002) Spatial and seasonal variations in stem respiration of beech trees (Fagus sylvatica). Annals of Forest Science 59, 801–812.

Chan T., Berninger F., Kolari P., Nikinmaa E. & Hölttä T. (2018) Linking stem growth respiration to the seasonal course of stem growth and GPP of Scots pine. Tree Physiology 38, 1356–1370.

Chiti T., Papale D., Smith P., Dalmonech D., Matteucci G., Yeluripati J., … Valentini R. (2010) Predicting changes in soil organic carbon in mediterranean and alpine forests during the Kyoto Protocol commitment periods using the CENTURY model. Soil Use and Management 26, 475–484.

Collalti A., Marconi S., Ibrom A., Trotta C., Anav A., D’Andrea E., … Santini M. (2016) Validation of 3D-CMCC Forest Ecosystem Model (v . 5 . 1) against eddy covariance data for 10 European forest sites. Geoscientific Model Development 9, 479–504.

Collalti A., Tjoelker M.G., Hoch G., Mäkelä A., Guidolotti G., Heskel M., … Prentice I.C. (2019) Plant respiration: Controlled by photosynthesis or biomass? Global Change Biology 0.

Collalti A., Trotta C., Keenan T.F., Ibrom A., Bond-lamberty B., Grote R., … Reyer C.P.O. (2018) Thinning Can Reduce Losses in Carbon Use Efficiency and Carbon Stocks in Managed Forests Under Warmer Climate. Journal of Advances in Modeling Earth Systems, 2427–2452.

Čufar K., Prislan P., De Luis M. & Gričar J. (2008) Tree-ring variation, wood formation and phenology of beech (Fagus sylvatica) from a representative site in Slovenia, SE Central Europe. Trees - Structure and Function 22, 749–758.

Cuny H.E., Rathgeber C.B.K., Frank D., Fonti P., Mäkinen H., Prislan P., … Fournier M. (2015) Woody biomass production lags stem-girth increase by over one month in coniferous forests. Nature Plants 1, 15160.

D’Andrea E., Rezaie N., Battistelli A., Kuhlmann I., Matteucci G., Moscatello S., … Muhr J. (2019) Winter’s bite□: beech trees survive complete defoliation due to spring late-frost damage by mobilizing old C reserves. 224, 625–631.

Damesin C., Ceschia E., Le Goff N., Ottorini J.-M. & Dufrêne E. (2002) Stem and branch respiration of beech: from tree measurements to estimations at the stand level. New Phytologist 153, 159–172.

Delpierre N., Lireux S., Hartig F., Camarero J.J., Cheaib A., Čufar K., … Rathgeber C.B.K. (2019) Chilling and forcing temperatures interact to predict the onset of wood formation in Northern Hemisphere conifers. Global Change Biology 25, 1089–1105.

Deslauriers A., Rossi S., Anfodillo T. & Saracino A. (2008) Cambial phenology, wood formation and temperature thresholds in two contrasting years at high altitude in southern Italy. Tree physiology 28, 863–871.

Dittmar C., Fricke W. & Elling W. (2006) Impact of late frost events on radial growth of common beech (Fagus sylvatica L .) in Southern Germany. European Journal of Forest Research 125, 249–259.

Di Filippo A., Biondi F., Maugeri M., Schirone B. & Piovesan G. (2012) Bioclimate and growth history affect beech lifespan in the Italian Alps and Apennines. Global Change Biology 18, 960–972.

Forner A., Valladares F., Bonal D., Granier A. & Grossiord C. (2018) Extreme droughts affecting Mediterranean tree species’ growth and water-use efficiency: the importance of timing. Tree physiology.

Frank D., Reichstein M., Bahn M., Thonicke K., Frank D., Mahecha M.D., … Zscheischler J. (2015) Effects of climate extremes on the terrestrial carbon cycle: Concepts, processes and potential future impacts. Global Change Biology 21, 2861–2880.

Gazol A., Camarero J.J., Colangelo M., de Luis M., Martínez del Castillo E. & Serra-Maluquer X. (2019) Summer drought and spring frost, but not their interaction, constrain European beech and Silver fir growth in their southern distribution limits. Agricultural and Forest Meteorology 278, 107695.

Gordo O. & Sanz J.J. (2010) Impact of climate change on plant phenology in Mediterranean ecosystems. Global Change Biology 16, 1082–1106.

Greco S., Infusino M., De Donato C., Coluzzi R., Imbrenda V., Lanfredi M., … Scalercio S. (2018) Late spring frost in mediterranean beech forests: Extended crown dieback and short-term effects on moth communities. Forests 9, 1–18.

Gričar J., Krže L. & Čufar K. (2009) Number of cells in xylem, phloem and dormant cambium in silver fir (ABIES ALBA), in trees of different vitality. IAWA Journal, 30, 121–133.

Gruber A., Wieser G. & Oberhuber W. (2009) Intra-annual dynamics of stem CO2 efflux in relation to cambial activity and xylem development in Pinus cembra. Tree Physiology 29, 641–649.

Guidolotti G., Rey A., D’Andrea E., Matteucci G. & De Angelis P. (2013) Effect of environmental variables and stand structure on ecosystem respiration components in a Mediterranean beech forest. Tree Physiology 33, 960–972.

Hilman B., Muhr J., Trumbore S.E., Kunert N., Carbone M.S., Yuval P., … Angert A. (2019) Comparison of CO2 and O2 fluxes demonstrate retention of respired CO2 in tree stems from a range of tree species. Biogeosciences 16, 177–191.

Katayama A., Kume T., Ichihashi R. & Nakagawa M. (2019) Vertical variation in wood CO2 efflux is not uniformly related to height: measurement across various species and sizes of Bornean tropical rainforest trees. Tree Physiology 39, 1000–1008.

Lempereur M., Martin-stpaul N.K., Damesin C., Joffre R., Ourcival J., Rocheteau A. & Rambal S. (2015) Growth duration is a better predictor of stem increment than carbon supply in a Mediterranean oak forest : implications for assessing forest productivity under climate change. 579–590.

Linares J.C., Camarero J.J. & Carreira J.A. (2009) Plastic responses of Abies pinsapo xylogenesis to drought and competition. Tree Physiology 29, 1525–1536.

Matteucci G. (1998) Bilancio del carbonio in una faggeta dell’Italia Centro-Meridionale: determinanti ecofisiologici, integrazione a livello di copertura e simulazione dell’impatto dei cambiamenti ambientali. (ed Università degli studi di Padova), Padova.

Meir P., Mencuccini M. & Coughlin S.I. (2019) Respiration in wood: integrating across tissues, functions and scales. New Phytologist, 1824–1827.

Merganičová K., Merganič J., Lehtonen A., Vacchiano G., Sever M.Z.O., Augustynczik A.L.D., … Reyer C.P.O. (2019) Forest carbon allocation modelling under climate change. Tree Physiology 39, 1937–1960.

Michelot A., Simard S., Rathgeber C., Dufrêne E. & Damesin C. (2012) Comparing the intra-annual wood formation of three European species (Fagus sylvatica, Quercus petraea and Pinus sylvestris) as related to leaf phenology and non-structural carbohydrate dynamics. Tree Physiology 32, 1033–1045.

Mund M., Kutsch W.L., Wirth C., Kahl T., Knohl a, Skomarkova M. V & Schulze E.-D. (2010) The influence of climate and fructification on the inter-annual variability of stem growth and net primary productivity in an old-growth, mixed beech forest. Tree physiology 30, 689–704.

Nolè A., Rita A., Ferrara A.M.S. & Borghetti M. (2018) Effects of a large-scale late spring frost on a beech (Fagus sylvatica L.) dominated Mediterranean mountain forest derived from the spatio-temporal variations of NDVI. Annals of Forest Science 75, 83.

Oladi R., Pourtahmasi K., Eckstein D. & Bräuning A. (2011) Seasonal dynamics of wood formation in Oriental beech (Fagus orientalis Lipsky) along an altitudinal gradient in the Hyrcanian forest, Iran. Trees - Structure and Function 25, 425–433.

Paembonan S.A., Hagihara A. & Hozumi K. (1992) Long-Term respiration in relation to growth and maintenance processes of the aboveground parts of a hinoki forest tree. Tree Physiology 10, 21–31.

Peuke A.D., Schraml C., Hartung W. & Rennenberg H. (2002) Identification of drought-sensitive beech ecotypes by physiological parameters. New Phytologist 154, 373–387.

Piovesan G., Biondi F., Di Filippo A., Alessandrini A. & Maugeri M. (2008) Drought-driven growth reduction in old beech (Fagus sylvatica L.) forests of the central Apennines, Italy. Global Change Biology 14, 1265–1281.

Principe A., Struwe T., Wilmking M. & Kreyling J. (2017) Low resistance but high resilience in growth of a major deciduous forest tree (Fagus sylvatica L.) in response to late spring frost in southern Germany. Trees - Structure and Function, 743–751.

Prislan P., Čufar K., De Luis M. & Gričar J. (2018) Precipitation is not limiting for xylem formation dynamics and vessel development in European beech from two temperate forest sites. Tree Physiology 38, 186–197.

Prislan P., Gričar J., De Luis M., Smith K.T. & Cufar K. (2013) Phenological variation in xylem and phloem formation in Fagus sylvatica from two contrasting sites. Agricultural and Forest Meteorology, 142–151.

R Development Core Team (2018) R: A Language and Environment for Statistical Computing.

Rathgeber C.B.K., Rossi S. & Bontemps J.-D. (2011) Cambial activity related to tree size in a mature silver-fir plantation. Annals of Botany 108, 429–438.

Rathgeber C.B.K., Santenoise P. & Cuny H.E. (2018) CAVIAR: An R package for checking, displaying and processing wood-formation-monitoring data. Tree Physiology 38, 1246–1260.

Reyer C.P., Silveyra Gonzalez R., Dolos K., Hartig F., Hauf Y., Noack M., … Frieler K. (2019) The PROFOUND database for evaluating vegetation models and simulating climate impacts on forests. Earth System Science Data Discussion, 1–47.

Rezaie N., D’Andrea E., Bräuning A., Matteucci G., Bombi P. & Lauteri M. (2018) Do atmospheric CO2 concentration increase, climate and forest management affect iWUE of common beech? Evidences from carbon isotope analyses in tree rings. Tree Physiology 1975, 1110–1126.

Rita A., Camarero J.J., Nolè A., Borghetti M., Brunetti M., Pergola N., … Ripullone F. (2019) The impact of drought spells on forests depends on site conditions□: The case of 2017 summer heat wave in southern Europe. 1–13.

Rossi S., Deslauriers A., Anfodillo T., Morin H., Saracino A., Motta R. & Borghetti M. (2006) Conifers in cold environments synchronize maximum growth rate of tree-ring formation with day length. New Phytologist 170, 301–310.

Rossi S., Menardi R., Fontanella F. & Anfodillo T. (2005) Campionatore Trephor: un nuovo strumento per l’analisi della xilogenesi in specie legnose. Dendronatura 1, 60–67.

Saveyn A., Steppe K., Mc Guire M.A., Lemeur R. & Teskey Æ.R.O. (2008) Stem respiration and carbon dioxide efflux of young Populus deltoides trees in relation to temperature and xylem carbon dioxide concentration. Oecologia 154, 637–649.

Scarascia-Mugnozza G., Bauer G.A., Persson H., Matteucci G. & Masci A. (2000) Tree Biomass, Growth and Nutrient Pools. In Carbon and Nitrogen Cycling in European Forest Ecosystems, Ecological. (ed E.-D. Schulze), pp. 49–62. Springer Verlag.

Scartazza A., Moscatello S., Matteucci G., Battistelli A. & Brugnoli E. (2013) Seasonal and inter-annual dynamics of growth, non-structural carbohydrates and C stable isotopes in a Mediterranean beech forest. Tree physiology 33, 730–42.

Schröter D., Cramer W., Leemans R., Prentice I.C., Araújo M.B., Arnell N.W., … Zierl B. (2005) Ecosystem service supply and vulnerability to global change in Europe. Science (New York, N.Y.) 310, 1333–7.

Schume H., Grabner M. & Eckmüllner O. (2004) The influence of an altered groundwater regime on vessel properties of hybrid poplar. Trees 18, 184–194.

Stockfors J. & Linder S. (1998) Effect of nitrogen on the seasonal course of growth and maintenance respiration in stems of Norway spruce trees. Tree Physiology 18, 155–166.

Teskey R.O., Saveyn A., Steppe K. & McGuire M.A. (2008) Origin, fate and significance of CO_2_ in tree stems. New Phytologist 177, 17–32.

Trumbore S.E., Angert A., Kunert N., Muhr J. & Chambers J.Q. (2013) What’s the flux? Unraveling how CO2 fluxes from trees reflect underlying physiological processes. New Phytologist 197, 353–355.

Ubierna N., Kumar A.S., Cernusak L.A., Pangle R.E., Gag P.J. & Marshall J.D. (2009) Storage and transpiration have negligible effects on δ13C of stem CO2 efflux in large conifer trees. Tree Physiology 29, 1563–1574.

Vicente-Serrano S.M., Gouveia C., Camarero J.J., Beguería S., Trigo R., López-Moreno J.I., … Sanchez-Lorenzo A. (2013) Response of vegetation to drought time-scales across global land biomes. Proceedings of the National Academy of Sciences of the United States of America 110, 52–57.

Vitasse Y., Schneider L., Rixen C., Christen D. & Rebetez M. (2018) Increase in the risk of exposure of forest and fruit trees to spring frosts at higher elevations in Switzerland over the last four decades. Agricultural and Forest Meteorology 248, 60–69.

Yang J., He Y., Aubrey D.P., Zhuang Q. & Teskey R.O. (2016) Global patterns and predictors of stem CO 2 efflux in forest ecosystems. Global Change Biology 22, 1433– 1444.

Zhang X., Friedl M. a., Schaaf C.B., Strahler A.H., Hodges J.C.F., Gao F., … Huete A. (2003) Monitoring vegetation phenology using MODIS. Remote Sensing of Environment 84, 471–475.

