## supporting information for "Frost and drought: effects of extreme weather events on stem carbon dynamics in a Mediterranean beech forest" (from another version of this preprint: version 1, 2019-10-16)

Lateral surface allometric equation for Selva Piana beech trees

Thirty four trees, with the diameter at breast height (DBH) ranging from 12 - 40 cm (Masci,1995) representing the population of Selva Piana, were used for calculation of allometric equation to estimate stem lateral surface (Tab.1). The sampled trees of were felled down and stem were divided logs, whose the exact length, upper and lower diameter were measured.

For each log, lateral area (LA) was calculated according to truncated cone formula

$$LA=\pi(r_{1}+r_{2})\times\sqrt{(r_{1}-r_{2})^{2}+h^{2}}$$

Where r1, r2, and h are the lower and upper radius, and length of the log, respectively.

The lateral area of the stem of each sampled trees was the sum of all logs.

Hence we created a relationships among DBH and LA (Fig. 1), and tested graphically the four assumption of linear model ( homoscedasticity, normality of the error distribution, statistical independence of the errors, absence of influent points) (Fig. 2).

Table 1: Characteristics of the sampled trees used to make the allometric equations named TREE#/##

| **Tree_Id** | **DBH (cm)** | **Stem length(m)** | **Lat_area (m^2^)** |
| --- | --- | --- | --- |
| Tree 1/92 | 29 | 16.9 | 9.3 |
| Tree 2/92 | 40 | 21.4 | 16.6 |
| Tree 3/92 | 28 | 23 | 14.7 |
| Tree 4/92 | 32 | 23.9 | 16.2 |
| Tree 5/92 | 32.5 | 23.1 | 14.9 |
| Tree 6/92 | 14 | 17.8 | 5.1 |
| Tree 7/92 | 31 | 23.9 | 15.0 |
| Tree 8/92 | 24 | 19.2 | 10.1 |
| Tree 9/92 | 19 | 17.8 | 7.0 |
| Tree 10/92 | 22 | 19.7 | 8.7 |
| Tree 11/92 | 25 | 19 | 9.2 |
| Tree 12/92 | 28 | 18.5 | 11.0 |
| Tree 1/95 | 25.6 | 17.2 | 8.3 |
| Tree 2/95 | 21 | 21.95 | 8.7 |
| Tree 3/95 | 17.6 | 17.22 | 5.9 |
| Tree 4/95 | 17 | 16.9 | 6.2 |
| Tree 5/95 | 38 | 21.9 | 15.7 |
| Tree 6/95 | 33.5 | 19.6 | 11.7 |
| Tree 8/96 | 38.5 | 22.85 | 16.0 |
| Tree 9/96 | 24 | 15.43 | 7.0 |
| Tree 10/96 | 29.5 | 20.3 | 12.4 |
| Tree 11/96 | 35 | 18.3 | 11.6 |
| Tree 12/96 | 26 | 15.55 | 8.0 |
| Tree 13/96 | 23.5 | 20.9 | 10.5 |
| Tree 14/96 | 29.6 | 17.7 | 9.5 |
| Tree 15/96 | 20.7 | 14.8 | 6.0 |
| Tree 16/96 | 18.6 | 15.8 | 5.9 |
| Tree 17/96 | 28.3 | 18.45 | 9.9 |
| Tree 18/96 | 35 | 21.07 | 13.3 |
| Tree 19/96 | 21 | 19.16 | 7.9 |
| Tree 20/96 | 16 | 15.35 | 5.0 |
| Tree 21/96 | 32.4 | 19.85 | 13.4 |
| Tree 22/96 | 11.8 | 15.3 | 3.3 |

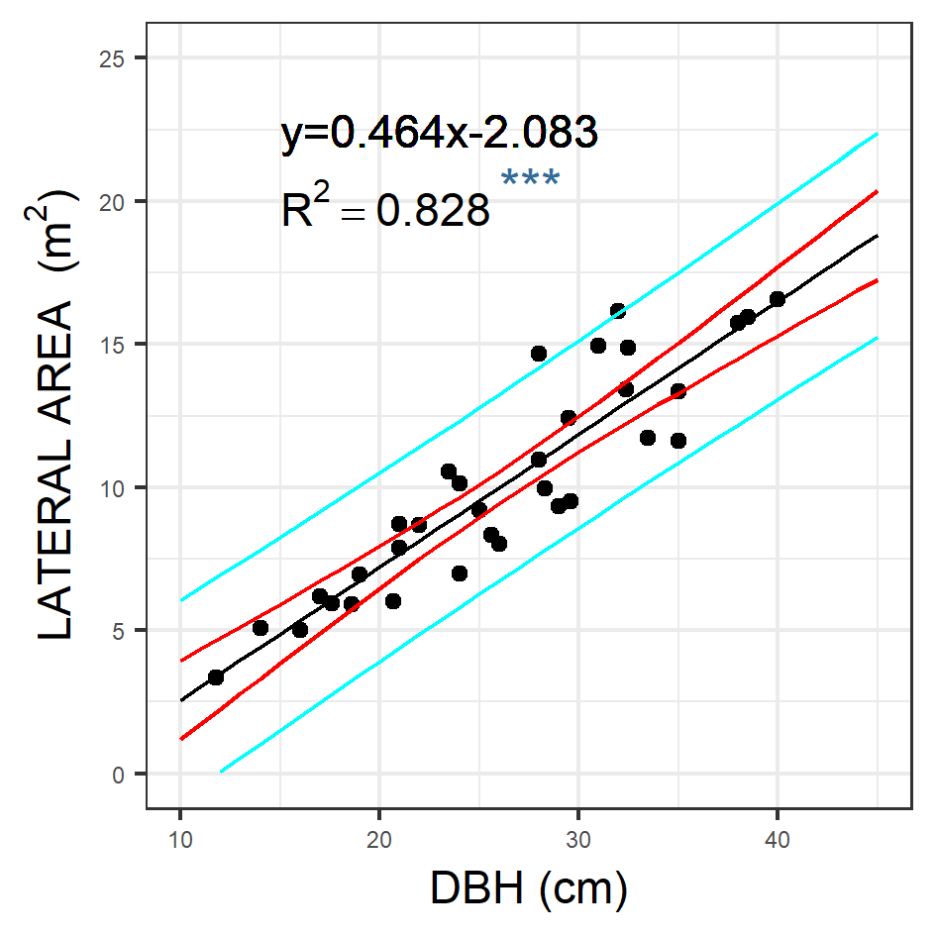

Figure 1: Allometric equation for calculation of beech stem Lateral Area. Each point is a Tree. Red lines are 95% Confidence interval. Cyan lines are prediction bands.

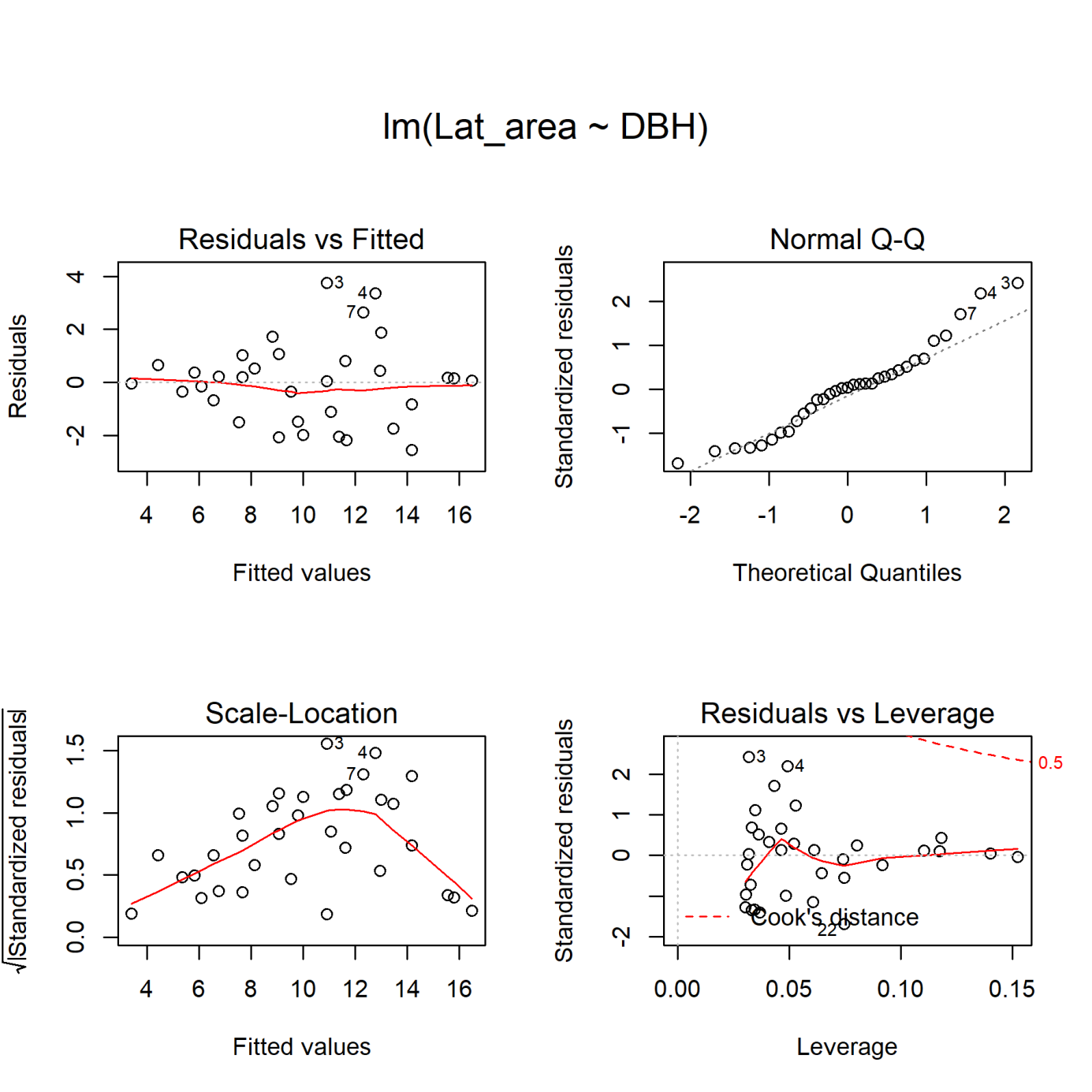

Figure 2: Graphical validation of the assumptions of linear model (homoscedasticity, normality of the error distribution, statistical independence of the errors, absence of influent points).

Relationships between ES and T air

Table 2: Parameters of ES vs Air temperature exponential relationship (y=ae^bx^) during non wood formation and wood formation periods , and relatives ES_15_ and Q_10_. Sampling dates and N represent the number of sampling and samples included in a period.

| **Year** | **Tree** | **Period** | **a** | **b** | **R^2^** | **P-value** | **Sampling dates (N)** | **ES_15w_** | **Q_10w_** |
| --- | --- | --- | --- | --- | --- | --- | --- | --- | --- |
| 2015 | 1 | Wood formation | 0.213 | 0.116 | 0.681 | 0.006 | 10 (20) | 1.214 | 3.191 |
| 2015 | 2 |  | 0.212 | 0.125 | 0.691 | 0.002 | 11 (21) | 1.378 | 3.488 |
| 2015 | 3 |  | 0.128 | 0.119 | 0.703 | 0.001 | 10 (18) | 0.762 | 3.292 |
| 2015 | 4 |  | 0.079 | 0.133 | 0.719 | 0.004 | 9 (18) | 0.578 | 3.780 |
| 2015 | 5 |  | 0.226 | 0.112 | 0.640 | 0.003 | 11 (22) | 1.208 | 3.053 |
| 2016 | 1 |  | 0.194 | 0.080 | 0.633 | 0.058 | 6 (12) | 0.647 | 2.232 |
| 2016 | 2 |  | 0.107 | 0.125 | 0.834 | 0.011 | 6 (12) | 0.700 | 3.490 |
| 2016 | 3 |  | 0.178 | 0.076 | 0.819 | 0.013 | 6 (11) | 0.557 | 2.138 |
| 2016 | 4 |  |  |  |  |  |  |  |  |
| 2016 | 5 |  | 0.365 | 0.051 | 0.799 | 0.003 | 8 (16) | 0.788 | 1.670 |
| 2017 | 1 |  | 0.102 | 0.118 | 0.780 | 0.041 | 6 (12) | 0.599 | 3.254 |
| 2017 | 2 |  | 0.266 | 0.113 | 0.733 | 0.051 | 6 (12) | 1.449 | 3.096 |
| 2017 | 3 |  | 0.133 | 0.131 | 0.721 | 0.054 | 6 (12) | 0.949 | 3.706 |
| 2017 | 4 |  | 0.131 | 0.114 | 0.696 | 0.060 | 6 (12) | 0.724 | 3.127 |
| 2017 | 5 |  | 0.128 | 0.121 | 0.761 | 0.045 | 6 (12) | 0.787 | 3.362 |
| 2015 | 1 | Non wood formation | 0.236 | 0.083 | 0.759 | 0.011 | 6 (10) | 0.819 | 2.291 |
| 2015 | 2 |  | 0.330 | 0.076 | 0.843 | 0.001 | 5 (9) | 1.032 | 2.206 |
| 2015 | 3 |  | 0.190 | 0.096 | 0.943 | 0.001 | 6 (10) | 0.802 | 2.094 |
| 2015 | 4 |  | 0.253 | 0.050 | 0.608 | 0.039 | 7 (13) | 0.536 | 1.651 |
| 2015 | 5 |  | 0.321 | 0.070 | 0.815 | 0.014 | 5 (9) | 0.917 | 2.014 |
| 2016 | 1 |  | 0.309 | 0.045 | 0.908 | 0.014 | 6 (12) | 0.607 | 1.568 |
| 2016 | 2 |  | 0.447 | 0.053 | 0.882 | 0.005 | 6 (12) | 0.983 | 1.692 |
| 2016 | 3 |  | 0.255 | 0.060 | 0.810 | 0.015 | 6 (11) | 0.628 | 1.822 |
| 2016 | 4 |  | 0.192 | 0.049 | 0.744 | 0.000 | 12 (24) | 0.404 | 1.639 |
| 2016 | 5 |  | 0.236 | 0.058 | 0.986 | 0.007 | 4 (8) | 0.559 | 1.777 |
| 2017 | 1 |  | 0.431 | 0.032 | 0.761 | 0.043 | 6 (12) | 0.697 | 1.377 |
| 2017 | 2 |  | 0.402 | 0.066 | 0.716 | 0.055 | 6 (12) | 1.082 | 1.935 |
| 2017 | 3 |  | 0.174 | 0.095 | 0.862 | 0.020 | 6 (12) | 0.724 | 2.586 |
| 2017 | 4 |  | 0.261 | 0.042 | 0.802 | 0.040 | 6 (12) | 0.490 | 1.522 |
| 2017 | 5 |  | 0.152 | 0.102 | 0.752 | 0.047 | 6 (12) | 0.706 | 2.783 |
| 2015 | 1 | Annual | 0.163 | 0.128 | 0.767 | 0.000 | 16 | 1.100 | 3.578 |
| 2015 | 2 |  | 0.227 | 0.122 | 0.798 | 0.000 | 16 | 1.405 | 3.369 |
| 2015 | 3 |  | 0.147 | 0.112 | 0.791 | 0.000 | 16 | 0.790 | 3.068 |
| 2015 | 4 |  | 0.100 | 0.120 | 0.727 | 0.000 | 16 | 0.604 | 3.322 |
| 2015 | 5 |  | 0.226 | 0.112 | 0.784 | 0.000 | 16 | 1.208 | 3.055 |
| 2016 | 1 |  | 0.320 | 0.052 | 0.500 | 0.010 | 12 | 0.694 | 1.674 |
| 2016 | 2 |  | 0.279 | 0.077 | 0.736 | 0.000 | 12 | 0.884 | 2.159 |
| 2016 | 3 |  | 0.224 | 0.065 | 0.798 | 0.000 | 12 | 0.596 | 1.924 |
| 2016 | 4 |  | 0.192 | 0.049 | 0.744 | 0.000 | 12 | 0.404 | 1.639 |
| 2016 | 5 |  | 0.297 | 0.059 | 0.787 | 0.000 | 12 | 0.717 | 1.801 |
| 2017 | 1 |  | 0.076 | 0.165 | 0.857 | 0.000 | 12 | 0.901 | 5.220 |
| 2017 | 2 |  | 0.258 | 0.113 | 0.765 | 0.000 | 12 | 1.402 | 3.095 |
| 2017 | 3 |  | 0.111 | 0.139 | 0.835 | 0.000 | 12 | 0.889 | 3.995 |
| 2017 | 4 |  | 0.094 | 0.129 | 0.817 | 0.000 | 12 | 0.649 | 3.618 |
| 2017 | 5 |  | 0.119 | 0.124 | 0.890 | 0.000 | 12 | 0.769 | 3.461 |

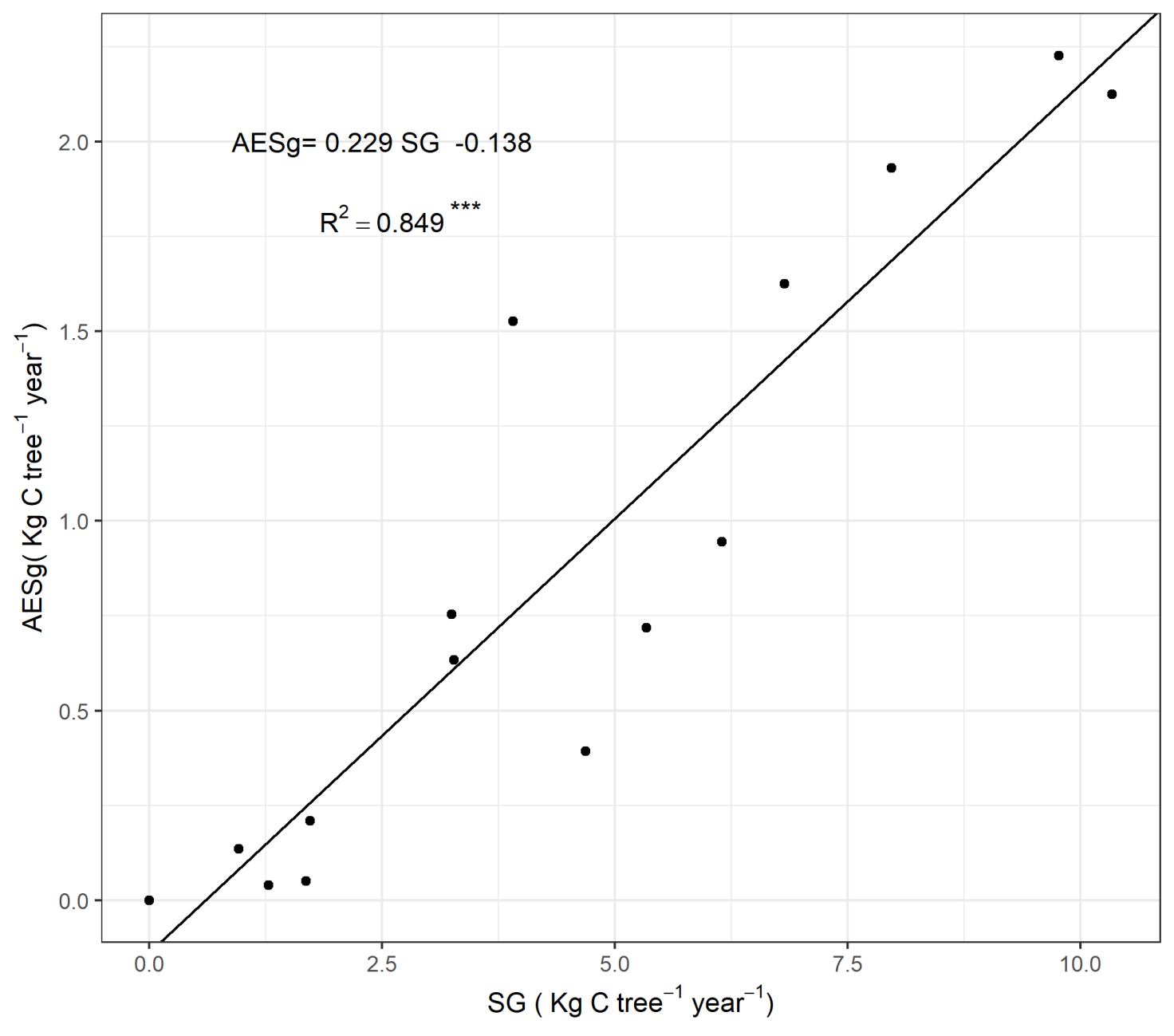
